# A sublethal drought and rewatering time course reveals intricate patterning of responses in the annual Arabidopsis thaliana

**DOI:** 10.1101/2025.07.25.666782

**Authors:** Elisabeth Fitzek-Campbell, Dennis Psaroudakis, Bernd Weisshaar, Astrid Junker, Andrea Bräutigam

**Affiliations:** University Bielefeld, Computational Biology, Faculty of Biology, CeBiTec, Universitätsstr. 27, 33615 Bielefeld, Germany; Klinikum Bielefeld, Institut for laboratory medicine, Microbiology and Transfusion medicine, Teutoburger Str. 50, 33604 Bielefeld; Leibniz Institute of Plant Genetics and Crop Plant Research (IPK), Germany; Forschungszentrum Jülich GmbH, CEPLAS, Bioeconomy Science Center (BioSC), Institute of Bio- and Geosciences (IBG-4: Bioinformatics), 52428 Jülich, Germany; Syngenta Seeds GmbH, Zum Knipkenbach 20, 32107, Bad Salzuflen, Germany

**Keywords:** progressive drought stress, recovery, ABA, anthocyanin pathway, TFs

## Abstract

Drought is a major factor of yield loss in annual crops. It triggers a wide range of physiological and molecular changes which are mediated by a suite of transcription factors from at least four different protein families. In the literature, the observed phenotypic changes and the number of transcriptome changes upon drought in *Arabidopsis thaliana* varies widely. To resolve the apparent variation, we conducted a phenotyping and transcriptomics experiment with progressive drought which is initially mild and escalates to strong but still sublethal drought followed by rewatering. Phenotypic data is analyzed with machine learning methods and connected to transcriptome data.

The phenotypic data show that drought stress manifests in distinct stages. The transcriptional analysis shows one threshold program and gradual expression programs among typical drought responsive regulons over time and during recovery. Plant aging prior to senescence and the drought response overlap to a large degree and drought stressed plants rejuvenate transcriptionally before returning to the control aging program. The phenotypic traits are associated with different transcript abundances again reflecting multiple overlaying programs. Transcripts with high explanatory power of phenotypes are biomarkers and not causal for the phenotype.

**Significance statement:** Large scale phenotyping and transcriptomics during progressive drought demonstrate that the drought response in the annual *Arabidopsis thaliana* is controlled by multiple transcriptional programs, mostly with gradual onset, that overlap with the plant aging program. Transcripts with high explanatory power for phenotypes are biomarkers and not causal.

## Introduction

Drought stress in annual plants causes substantial agricultural yield loss (Dietz et al., 2021). Reduced soil water content leads to drought stress which constrains plant growth, development, photosynthetic carbon fixation, and ultimately plant productivity (Dietz et al., 2021). Plants have evolved a complex system of mechanisms to manage drought stress which include changes at the morphological, physiological, biochemical, and regulatory level. Physiological and morphological changes include reduced leaf area and stem length as well as reduced growth rates (Matthews et al., 1984). Leaf water potential and the relative water content decrease (Lawlor and Cornic, 2002). Stomatal conductance is reduced (Cornic, 2000) and a reduced photosynthetic rate is observed (Lawson, 2003; Tezara et al., 1999). Photosystem II chemistry is altered during water stress (Lu and Zhang, 1999). Depending on the duration and severity of the stress, yield is reduced (Santini et al., 2022). On the biochemical level, reactive oxygen circuits are altered with changes in antioxidative enzymes and changes in reactive oxygen species (ROS) accumulation (summarized in (Reddy et al., 2004)). The enzymes involved in managing ROS include superoxide dismutase (SOD), ascorbate peroxidase (APX), monodehydroascorbate reductase (MDAR), peroxidase (POD), glutathione reductase (GR), alternative oxidase (AOX), and catalase (CAT). Also, metabolites such as flavanones, glutathiones, anthocyanins, and ascorbic acid mediate ROS detoxification (Apel and Hirt, 2004; Reddy et al., 2004). At the same time, chlorophyll levels are reduced (Mafakheri et al., 2010; Wu et al., 2016), and anthocyanins (Chalker-Scott, 1999), compatible solutes such as sugars, proline (Hare et al., 1998) as well as protective proteins such as LATE EMBRYO ABUNDANT (LEA) proteins accumulate (Hundertmark and Hincha, 2008). Drought responses of plants are versatile and a given species at a given developmental stage may respond with different mechanisms (Fang and Xiong, 2015). At the regulatory level, the plant hormone abscisic acid (ABA) mediates a large proportion of the drought response which is triggered by substantial changes at the level of transcription (Shinozaki and Yamaguchi-Shinozaki, 2006; Tenorio Berrío et al., 2022). Together with ABA, the phytohormones ethylene, jasmonic acid, and cytokinin orchestrate a cross-talk in the response to abiotic stresses such as osmotic, drought, heat, cold, salt and UV radiation (Cutler et al., 2010; Divi et al., 2010; Finkelstein, 2013, 2013; Gazzarrini and Mccourt, n.d.; Ruan et al., 2010).

ABA is synthesized from zeaxanthin in chloroplasts and the cytosol (Finkelstein, 2013). The activities of the ABA biosynthetic enzymes ZEP, NCED4, AAO3 and ABA3 are rapidly upregulated after dehydration stress (3, 6, 9h). The late biosynthetic enzyme ABA2 functions as a late-responsive gene and is responsible for fine-tuning the biosynthesis of ABA (Lin et al., 2007). ABA is recognized by Pyrabactin Resistant/Pyrabactin Resistant-Like/Regulatory Component of ABA Response (PYR/PYL/RCAR) receptors (Muhammad Aslam et al., 2022; Yoshida et al., 2014). Upon drought stress, endogenous ABA accumulates in the cell and binds to its receptor which induces a conformational change that releases protein phosphatases 2C (PP2C) to transduce the signal by phosphorylating SNF1-Related Protein Kinase 2 (SnRK2) (Finkelstein, 2013). SnRK2 acts on transcription factors (TFs).

Within the gene regulatory networks, multiple regulons respond to drought (Yamaguchi-Shinozaki and Shinozaki, 2006). The AREB/ABF TFs, part of the bZIP family, respond to increases in endogenous ABA and bind the ABA responsive element (ABRE) in their target gene promoters to affect the drought response (Choi et al., 2000; Kulik et al., 2011; Umezawa et al., 2009). The DREB/CBF TFs of the ERF/AP2 family respond independently of ABA and bind the DRE/CRT element in their target promoters to affect the drought response (Agarwal et al., 2017; Lata and Prasad, 2011; Yamaguchi-Shinozaki and Shinozaki, 2006, 1993). Especially the DREB2 subfamily TFs respond to and mediate a drought response (Nakashima et al., 2000) while other family members are activated by other abiotic stresses such as cold and/or heat (Fujita et al., 2011). DREB/CBF TFs are at the cross-roads to different hormones (Akhtar et al., 2012). In the perennial *Talinum fruticosum* (formerly *Talinum triangulare*), they mediate an initial but not a prolonged response to drought (Brilhaus et al., 2016). Several other transcription factors such as NACs and WRKYs also take part in the drought response independent of ABA (Yamaguchi-Shinozaki and Shinozaki, 2006).

Anthocyanin accumulation is induced not only by drought, but also by high light, cold and salinity. Anthocyanins can function as sunscreen by scavenging excess ROS to prevent damage of photosynthetic tissue (Chalker-Scott, 1999; Kovinich et al., 2014; Li and Ahammed, 2023). The anthocyanin biosynthesis pathway is governed by a network of MYB TFs and bHLH TFs, respectively (Borevitz et al., 2000; Stracke et al., 2010). The overexpression of anthocyanin regulating genes (Nakabayashi et al., 2014), of ABFs (Choi et al., 2000) and of DRE/CBFs (Akhtar et al., 2012) confer some aspects of drought tolerance and increase plant resilience, underscoring their vital role in the drought response network.

A large number of transcriptome studies have determined drought responsive transcripts in *A. thaliana* resulting in very different numbers of regulated transcripts. Initial single time point expression profiling with 7000 transcripts identified 277 drought inducible transcripts with more than five-fold induction including upregulation of 40 TFs, of 11 osmoprotectant synthesis genes, and 5 LEA genes and downregulation of 37 photosynthesis genes (Seki et al., 2002). Early studies focused on harsh artificial shock treatments and few or a single time point (Abdeen et al., 2010; Bechtold et al., 2016; Deyholos, 2010; Fujita et al., 2009; Kawaguchi et al., 2004; Kilian et al., 2007; Kreps et al., 2002; Mizoguchi et al., 2010; Seki et al., 2002; Weston et al., 2008). A meta-analysis of 11984 transcripts on microarrays revealed 4015 transcripts or about a third of all tested transcripts as differentially abundant across drought stress experiments including those in the ABA response (Rest et al., 2016). Later studies analyzed soil-grown plants under moderate drought conditions induced by water withholding and the analyses of time courses rather than single time points (Bechtold et al., 2016; Brilhaus et al., 2016; Harb et al., 2010; Wilkins et al., 2010). Drought treatment for 8 and 13 days, respectively, showed ABA responses early and acclimation via growth reduction late (Harb et al., 2010). A drought treatment for three days showed growth changes already and identified 759 probe sets sampled at four time points after prolonged drought as responding to drought with substantial fluctuation during the diurnal cycle (Wilkins et al., 2010). Treatment at day 35 after sowing with daily sampling for 14 days identified a total of 1815 differentially abundant transcripts (Bechtold et al., 2016). In a perennial plant, many, but not all transcripts at least partially recovered during rewatering and a model proposed a transient general stress signal and a sustained ABA mediated change which mostly reverted upon rewatering (Brilhaus et al., 2016). Taken together these analyses and the meta-analysis suggest that the drought response is highly variable depending on severity of drought, sampling time point, and method of application but contains a common core of mostly ABA responsive genes.

Given the large variation between experiments, we hypothesized that multiple overlying programs operate during the drought response. A time course from early mild drought towards strong, but sublethal drought at 10% field capacity in soil with both phenotype and transcriptome data was generated. The data showed that at least two transcriptional programs are operating. Increasing drought over time increased the transcriptional response. Control plants revealed a transcriptional aging program which was sped up in drought treated plants and reversed upon watering. The comparative analyses of phenotype and transcriptome data demonstrated that, with the exception of anthocyanin biosynthesis, the apparent threshold onset of phenotypic responses is mediated by gradual transcriptional changes in target genes and a large assortment of transcription factors.

## Material and methods

### Plant material and growth conditions

Seeds of *Arabidopsis thaliana* Col-0 underwent imbibition on wet filter paper overnight at 20° C and no light and were then transferred to soil (85% (v) red substrate 1 (Klasmann-Deilmann GmbH, Geeste, Germany) and 15% (v) sand, pH 5.5). A total of 137 pots (10 cm diameter, 8 cm height) were watered with 50 ml water each, covered with plastic caps to maintain high humidity, and consequently subjected to stratification conditions for two days in the dark at 5° C and 75 % relative humidity. To initiate germination, pots were exposed to a 16/8 h day/night cycle at 16/14° C, 75% relative humidity, and 120 μmol light intensity for one day (Whitelux Plus metal halide lamps (Venture Lighting Europe Ltd., Rickmansworth, Hertfordshire, England)), then light intensity was increased to 180 μmol for two days. After germination, plastic caps were removed and conditions stayed stable at 20/18° C and 60/75 % humidity for a 16/8 h day/night rhythm with 180 μmol light during the day.

Soil water level was determined through field capacity at an automatic weighing and watering station the plants/pots visited once a day. 100 % field capacity corresponded to the weight of fully watered soil, 0 % capacity weight was determined by drying soil for 3 days at 80° C. Whenever plants arrived at the watering station, target weight for 80 % field capacity was calculated and the weight difference was added in water for the well-watered group(ignoring plant biomass weight).

Prior to the prolonged drought stress treatment, plants were watered and phenotyped. Starting at 15 DAS (days after sawing) after phenotyping and a final watering, 98 of the plants were subjected to drought stress by not being watered until DAS 30, the other 39 plants remained well-watered. On DAS 31, baskets were installed to stabilize the now flowering plants. Because the basket covers parts of the plant and can thereby interfere with measurements, 11 plants were chosen of which the flower shoot was regularly removed to generate some data that are unaltered by basket masking (Plants 140, 175, 226, 254, 269, 273, 277, 287, 299, 316, and 384). 44 days after sowing all remaining plants were harvested (cutting directly above the ground, without roots) and their fresh and dry weight was measured (Denver Instruments MXX-123 with 0.001 g precision, drying occurred in a drying oven at 80° C for 3 days).

### Phenotyping and Image Analysis

Starting at DAS 7, plants were imaged from top view with fluorescent light (FLUO, excitation: 400-500 nm, emission: 520–750 nm) using a Basler Scout scA1400-17gc (RGB) camera with a resolution of 1624×1234 pixels. On DAS 12, daily imaging with visible light was added to the routine (∼390–750 nm, Basler (Basler AG, Ahrensburg, Germany) Pilot piA2400-17gc (RGB) camera with a resolution of 2454×2056 pixels). Beginning with DAS 20, the zoom level configuration was switched to better capture the now bigger plants.

Images were automatically analyzed using the Integrated Analysis Platform (IAP) (Klukas et al., 2014). Details on the used IAP configuration can be found in (Arend et al., 2016). This yielded 173 phenotypic traits out of which we selected 42 biologically interpretable and non-equivalent traits (Supplementary Table S1).

### Count of publicly available transcriptome studies

To get an overview of how many transcriptome studies have been published and are publicly available, a search among Gene Expression Omnibus datasets with the search term “((drought) AND leaf) AND arabidopsis thaliana[Organism]” was conducted. The search recovered 610 datasets (retrieved 2024-03-19).

### RNA-sequencing

Leaf material of plants was sampled at noon on DAS 15 (before onset of drought), 17, 20, 22 (including control), 24, 27, 29 (including control), 31, 34, and 36 (including control) in three replicates. Libraries were prepared with TruSeq RNA Sample Prep Kit v2, quantified with Qubit 2.0 (Invitrogen) and single-end sequenced to an average depth of 12 million reads with 150 bases using a NextSeq500. Reads were mapped with kallisto v0.44.0 (with default parameters including the single end read flag --single -l 200 -s 30) on the TAIR10 transcriptome reference (Berardini et al., 2015; Bray et al., 2016). Principal component analysis was performed with log2 transformed, z-scaled data with prcomp in R. Differentially abundant transcripts compared to day 15 were determined with edgeR in classic mode followed by Benjamini-Hochberg multiple hypothesis correction (Robinson et al., 2010). Cluster analysis was performed with transcripts significantly differentially abundant on at least one time point with a partition around medoids clustering algorithm (PAM, cluster in R, with default parameters) from the cluster package in R (Rousseeuw et al., 2019). Gene ontology (GO) term enrichment analysis was performed with TopGO (default parameters except nodesize=10) from the Bioconductor package using ATH_GO_GOSLIM (release date 2023-04-01) obtained from TAIR (Alexa and Rahnenfuhrer, 2019). Data was visualized in R using UpsetR and ggplot2.

### Feature Selection

To identify which phenotypic traits are relevant to distinguish drought-stressed from well-watered plants, feature selection was performed in R (version 4.3.2 (R CoreTeam, 2021)) using the Boruta package (version 8.0.0, default parameters with maximum runs set to 1500 (Kursa and Rudnicki, 2010)) on days with complete and clean data (20 -35 DAS). Outliers were defined as being more than 2 standard deviations away from the median of that particular treatment group and DAS and, since random forest algorithms do not allow missing data, replaced by the median of the remaining data. 42 primary traits (Supplementary Table S1) were used as input. Results were categorized using the *TentativeRoughFix* function.

To identify genes with relevance for the prediction of each trait only transcripts showing differential abundance compared to DAS 15 (before the start of drought stress) were used and the required p-value cutoff was reduced to 0.01. Feature selection was performed using 1000 trees in 800 runs or until convergence. To further increase confidence in the result, the gene selection process was repeated and only the intersect set of all runs was kept. Iteration stopped when the resulting set did not get any smaller.

### Knockout Line Validation Experiment

To test the selected genes for causality, they were ranked using different criteria (number of associated traits and subnetworks, strength of association) to cover different scenarios and the overall top 230 ranking genes were screened for the existence of homozygous knockout mutants in the GABI-Kat and SALK collections (Supplementary Table S2). 71 knockout mutants covering 31 genes were then ordered and propagated along the background genotype Col0 to obtain seeds grown under identical conditions. DNA was extracted from the leaf tissue of the parental plants and homozygous insertion confirmed via PCR (Primer sequences in Supplementary Table S3). Seeds from 22 confirmed plants covering 11 genes with two independent knockout alleles per gene were then subjected to the same experimental and measurement protocols as above alongside the wildtype obtained from the same propagation and tested for a phenotype in drought conditions.

## Results

### Phenotypic characterization of sublethal drought

To accurately quantify the morphological changes in *Arabidopsis thaliana* in response to progressive drought stress, 137 plants of genotype Col0 were grown in a climate-controlled phyto chamber over the course of 44 days and measured daily using image-based, high-throughput phenotyping. Starting at 15 days after sowing (DAS), 98 plants were subjected to progressive, non-lethal drought stress by not being watered until 30 DAS (days after phenotyping), while 39 plants were kept well-watered throughout (Figure 1A). Phenotyping yielded a total of 545498 values covering 21 architectural and 18 color-related traits of which 12 were from the visible light spectrum, and 6 from the fluorescent light spectrum as well as the mean fluorescence and, starting at 20 DAS, the near-infrared (NIR) emission intensity. Growth in the drought treatment group was indistinguishable from control from DAS 15 to DAS 26, then slowed and stopped on DAS 29 as the water content in the soil decreased down to 10% field capacity (Figure 1B). For the days where complete data were available prior to basket installation to contain flowering shoots (DAS 20 to 35), the feature selection was applied to classify plants into either drought stress or well-watered treatment to determine which phenotypic traits are affected by the stress and at which time points. This reduced the feature spectrum of the 630 phenotypic input parameters to 67 relevant trait-day combinations by which we could classify the plants into their respective treatments with 100% accuracy in a leave-one-out cross-validation. The importance of traits increased with increasing drought severity until a maximum at 30 DAS when plants were rewatered (Figure 1C). On 14 DAS, only near infrared traits representing tissue water content are important for successful classification (Figure 1C, D). On 25 and 26 DAS or ten and eleven days of increasing drought near infrared visible traits gain importance (Figure 1C, E). Fluorescence based traits appear on day 27 while architectural traits only play a role in prediction from 29 DAS onwards when plants are facing severe, but sublethal drought (Figure 1C, F, G). At 31 DAS, after plants have been rewatered for 1 day the algorithm still finds traits of near infrared, visible and architecture types useful for prediction while on day 32 there are only visible traits of very minor importance (Figure 1C). The different times of onset for traits of the different categories may be due to different transcriptional regulation programs for the traits or different thresholds for each trait in the same transcriptional program.

**Figure 1.**
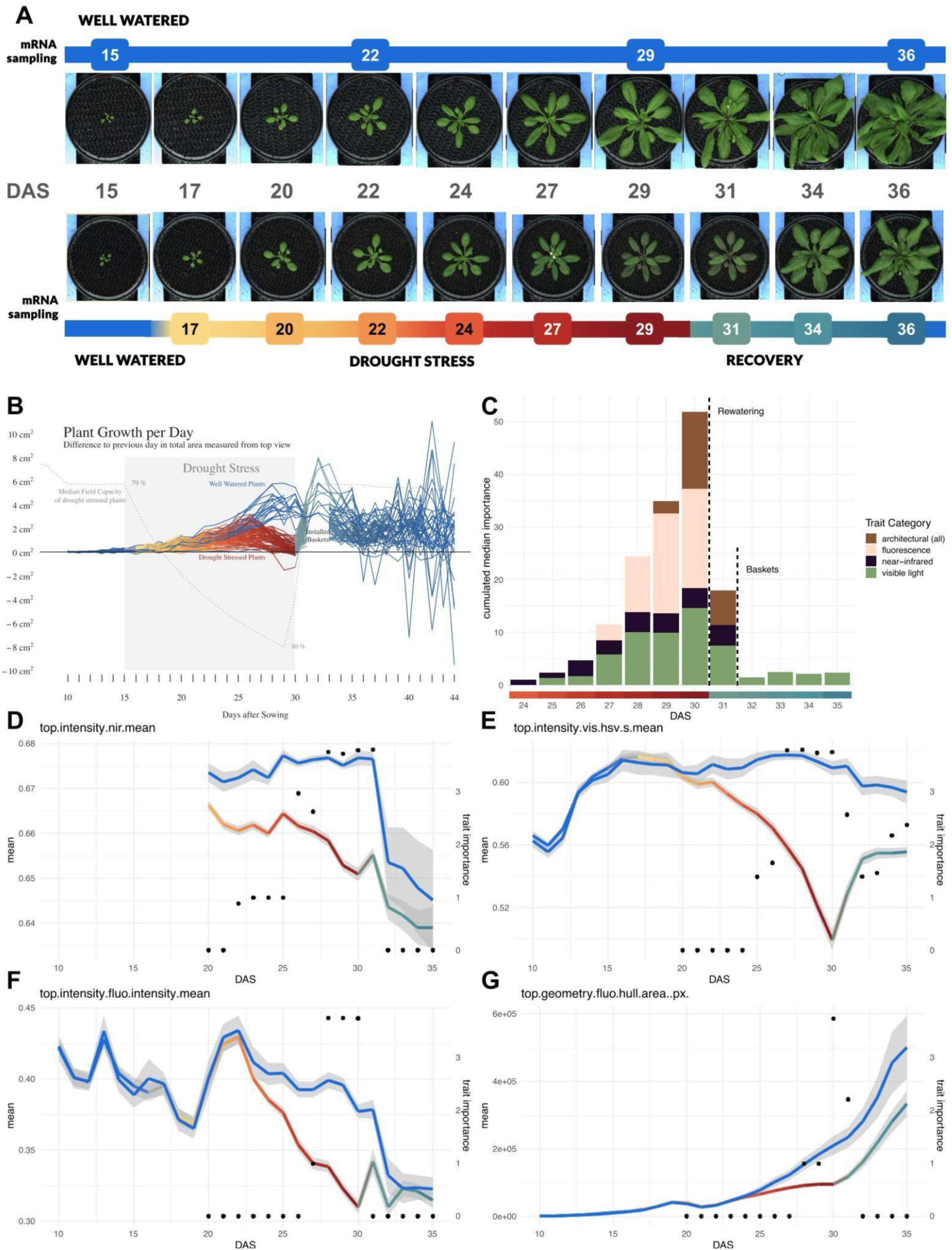
Phenotypes of progressive drought stress. **A**) Images of a representative drought-treatment and control plant throughout the experiment. For drought-treatment plants, watering was stopped after 15 days after sowing (DAS) and resumed on day 30. Leaf material was sampled for mRNA sequencing at the marked time points for the respective treatment groups. Brightness of images of 20 DAS and later was slightly increased to account for a shift in camera configuration. **B**) Absolute growth curves for all plants calculated as difference in leaf area compared to the previous day with variation after day 32 introduced by basket installation for flowering shoots. **C**) Importance of phenes (phenotypic traits) over time for distinguishing drought-stressed from well-watered plants assigned by the Boruta feature selection algorithm, cumulated by trait category. **D-G**) An exemplary trait from the categories near infrared (nir), visual (vis), fluorescence (fluo) and architecture (geometry) with control growth in blue and treatment drought and rewatering in color gradient with y-axis on the left and importance for prediction as black dots with y-axis on the right.

### Transcriptomic profiles of ABA signaling and anthocyanin biosynthesis show different patterns

To analyze the programs underlying the phenotypic symptoms of progressive drought stress we sampled leaf tissue for mRNA sequencing at 9 time points during drought stress and recovery and 4 time points for control plants (Figure 1A). To test the patterns of two well characterized signaling pathways relevant for drought, the pathway steps and controlling transcription factor abundance patterns were plotted next to their expression patterns (Figure 2). Overall, ABA synthesis and signaling transcripts increase over time with plant age of the control plants (Figure 2A, black dots) and they increase with progressive drought, drop directly upon rewatering, and increase again with age (Figure 2A, colored bars). This pattern holds for biosynthesis genes except for the beta carotene to zeaxanthin conversion by BCH genes and the xanthoxin to abscisic aldehyde conversion by ABA2. Among the signaling transcripts this pattern holds for RCAR2/PYL7 and RCAR1/PYL9 transcripts. Of the other isoforms of the ABA receptor, PYR1, PYL3 and PYL10 show an upregulation at day 29, whereas PYL1, PYL4, PYL5, PYL6, PYL8 and PYL11 show downregulation (Supplementary Figure 1). The three SnRKs, SnRK2.1, 2.4, and 2.6, also show the pattern of increase with progressive drought, drop upon rewatering, and increase with age (Figure 2A). Members of all SnRK subclasses show in general an upregulation at day 29 (Supplementary Figure 1). The six PP2Cs known to be involved in ABA signaling also show the typical pattern (Figure 2A). The seven bZIP transcription factors of group A known to be involved in ABA signaling show the typical pattern (Figure 2A). All members of group A bZIP TFs show highest upregulation at day 29 of the progressive drought stress, with the exception to DPBF2 that shows downregulation at the end of the drought stress treatment (Figure 2A, Supplementary Figure 1). Among the TFs with this typical response, the degree of response to rewatering varies with, for example, ABF3 and AREB3 falling way below levels detected on 15 DAS before the onset of stress while EEL drops to the level detected on 15 DAS (Figure 2A).

**Figure 2.**
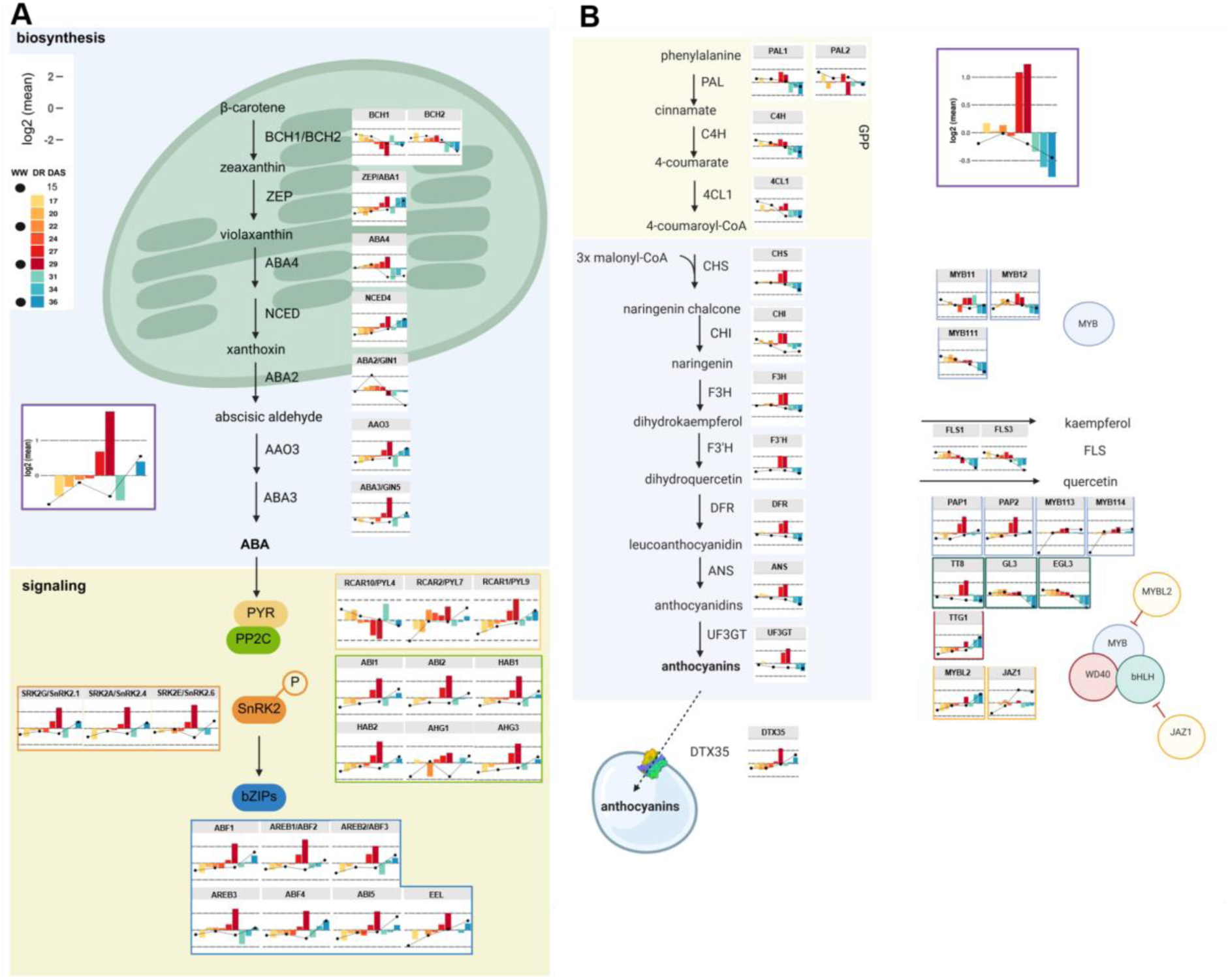
Transcriptome profiles of ABA synthesis and signaling pathway **(A)** and anthocyanin biosynthesis and their regulating TFs **(B).** The profiles show z-scores of log2 mean values relative to 15 DAS. Progressive drought and rewatering in colored bars and aging plants in control conditions in black points. The progressive drought stress is color coded from yellow-orange-red color scheme bars (one color per day), the intensity of the color is in correlation to the intensity of the progressive drought stress treatment. Rewatered treatment series is colored in green-blue bars (one color per day). The well-watered control was superimposed with connected black dots. The average transcriptional profile for both pathways is shown in purple box. Abbreviations: BETA CAROTENOID HYDROXYLASE 1 (BCH1), BETA CAROTENOID HYDROXYLASE 3 (BCH2), ABA DEFICIENT 1 (ABA1), NINE-CIS-EPOXYCAROTENOID DIOXYGENASE 4 (NCED4), ABA-deficient 4 (ABA4), ABA DEFICIENT 2 (ABA2), Abscisic ALDEHYDE OXIDASE 3 (AAO3), ABA DEFICIENT 3(ABA 3), RCAR (regulatory components of ABA receptor)10/ PYL(PYR1-like)4 (RCAR10/PYL4), RCAR (regulatory components of ABA receptor)2/ PYL(PYR1-like)7 (RCAR2/PYL7), RCAR (regulatory components of ABA receptor)1/ PYL(PYR1-like)9 (PCAR1/PYL9), ABA INSENSITIVE 1 (ABI1), ABA INSENSITIVE 2 (ABI2), HYPERSENSITIVE TO ABA1 (HAB1), HYPERSENSITIVE TO ABA2 (HAB2), ABA-hypersensitive germination 1 (AHG1), ABA-hypersensitive germination 3 (AHG3), SERINE/THREONINE KINASE 2G/SNF1-RELATED PROTEIN KINASE 2.1 (SRK2G/SnRK2.1), SERINE/THREONINE KINASE 2A/SNF1-RELATED PROTEIN KINASE 2.4 (SRK2A/SnRK2.4), SERINE/THREONINE KINASE 2E/SNF1-RELATED PROTEIN KINASE 2.6 (SRK2E/SnRK2.6), ABSCISIC ACID RESPONSIVE ELEMENT-BINDING FACTOR 1 (ABF1), ABSCISIC ACID RESPONSIVE ELEMENT-BINDING FACTOR 2 (ABF2), ABSCISIC ACID RESPONSIVE ELEMENT-BINDING FACTOR 3 (ABF3), ABA-RESPONSIVE ELEMENT BINDING PROTEIN 3 (AREB3), ABSCISIC ACID RESPONSIVE ELEMENT-BINDING FACTOR 4 (ABF4), ABA INSENSITIVE 5 (ABI5), ENHANCED EM LEVEL (EEL). General phenylpropanoid pathway(GPP) consists of: phenylalanine ammonia-lyase 1 (PAL1), phenylalanine ammonia-lyase 2 (PAL2), cinnamate-4-hydroxylase (C4H), 4-coumarate:CoA ligase 1 (4CL1). The anthocyanin pathway consists of the following steps: chalcone synthase (CHS), chalcone isomerase (CHI), flavanone 3-hydroxylase (F3H), flavonoid 3’ hydroxylase (F3’H), flavonol synthase 1&3 (FLS1 & 3), dihyroflavonol 4-reductase (DFR), anthocyanidin synthase (ANS), UDP-glucose flavonoid 3-O-glucsyltransferase (UF3GT), MATE efflux family protein (DTX35). TFs regulating the anthocyanin pathway: myb domain protein 11 (MYB11), myb domain protein 12 (MYB12), myb domain protein 111 (MYB111), production of anthocyanin pigment 1 (PAP1), production of anthocyanin pigment 2 (PAP2), myb domain protein 113 (MYB113), myb domain protein 114 (MYB114), TRANSPARENT TESTA 8 (TT8), GLABROUS 3 (GL3), ENHANCER OF GLABRA 3 (EGL3), TRANSPARENT TESTA GLABRA 1 (TTG1), MYB-like 2 (MYBL2), JASMONATE-ZIM-DOMAIN PROTEIN 1(JAZ1). Figure was generated in BioRender.

Overall, the expression of transcripts encoding proteins involved in the biosynthesis of anthocyanin decrease with plant age in control conditions. In drought conditions, they are induced at 27 and 29 DAS of progressive drought and unlike ABA-related transcripts permanently reduced upon rewatering (Figure 2B). This pattern holds for the transcripts yielding enzymes that produce 4-coumaroyl-CoA and for all transcripts leading to produce anthocyanin (Figure 2B). The transporter for sequestration of anthocyanins to the vacuole, DTX35, in contrast, increases with plant age and with progressive drought. Of the entry enzymes in specialized metabolism, phenylalanine ammonia ligases (PALs), only PAL1 displays this pattern typical for transcripts related to anthocyanin biosynthesis, while PAL 2 is decreased again on 29 DAS (Figure 2B) and PAL 3 shows increased expression at the beginning of the recovery phase (Supplementary Figure 2). The flavonol synthase (FLS) 1 and FLS3, which produce flavonol followed by hydroxylation to quercetin respectively, drain substrates from anthocyanin synthesis show stable or reduced expression at day 29 (Figure 2B). The proposed single MYB transcription factors MYB11, MYB12, and MYB111 of anthocyanin biosynthesis do not have the patterns of their supposed targets with MYB12 coming closest (Figure 2B). The TF-complex consisting of a tripartite complex of a WD40, a bHLH, and a MYB TF exhibit a diverse expression profile. For the candidate MYB-type TFs, PAP1 and PAP2 exactly mirror the expression of the target genes. For the bHLH component, TT8 exactly mirrors expression of the targets while GL3 and EGL3 stay flat during progressive drought stress and decline with age in both rewatering and control scenarios (Figure 2B). The WD40 TTG1 has a completely different expression pattern with increase over time irrespective of drought or control conditions (Figure 2B). The known negative regulators MYBL2 and JAZ1 are not anti correlated with their supposed targets (Figure 2B).

The analysis of two known drought-responsive regulons, that of ABA signaling and that of anthocyanin biosynthesis, reveal two different programs. One increases with progressive drought and with age, and responds with a drop below baseline upon rewatering; the other program has a threshold response on 27 DAS, returns to baseline after rewatering and decreases with age.

### Clustering and PCA analysis reveal premature aging under drought

To analyze the transcriptome data for additional patterns, the transcripts which showed significantly different abundance in at least one sampled time point were partitioned around medoids with PAM clustering. Of the ten clusters, eight showed continuous increase or decrease with age (Figure 3, dark blue lines in background). Only cluster 9 enriched in the flavonoid metabolic process (p=0.01) contains transcripts stable under control conditions. Clusters 2, 3, 6, 7 and 10 with 7.575 transcripts decrease with age in control conditions and clusters 1, 4, 5, and 8 with 7.665 transcripts increase with age in control conditions (Figure 3). The color gradient-coded drought stress time course in cluster 1 and 4 displays transcripts which increase faster compared to control conditions with a large spike at 29 DAS and a dip towards baseline upon rewatering. In cluster 1 the level catches up to control which at 36 DAS reaches the same level as the drought stress spike with transcripts enriched in catabolic processes (p=4.1e-79; Figure 3A), response gene ontology terms, and senescence (Supplementary Table S4). In cluster 4, the age spike at 36 DAS does not reach the drought stress spike at 29 DAS with transcripts enriched in the regulation of macromolecule metabolic processes, especially RNA biosynthetic processes (p=3.9e-09; Figure 3A; Supplementary Table S7). In cluster 5 enriched in response to stimulus (p=3.9e-48) including responses to bacteria and fungi (Supplementary Table S8) the drought stress induced spike remains below the age-related increase (Figure 3A). Cluster 8 behaves atypically. It is enriched in defense (p=3.5e-115), peaks very early in the stress time course at 20 DAS, drops again and then tracks control conditions closely (Figure 3A). Among the clusters with transcripts downregulated under control conditions with age, the patterns are more diverse. Clusters 6, 7, and 10 are similar to clusters 1 and 4 with transcripts decreasing faster compared to control with a downspike at 29 DAS. Cluster 6 is enriched in translation (p=7.6e-166) and translation-related processes such as rRNA and ribosome biogenesis (Supplementary Table S9) and transcripts overshoot the controls upon rewatering before catching up to control at 36 DAS (Figure 3A). Decline upon age in control conditions is only very slight in cluster 7 enriched in photosynthesis (p=9.2e-22; Figure 3A), and related processes such as chlorophyll biosynthesis, overshoot is small if at all present, and transcripts again catch up to control at 36 DAS. Cluster 10 contains transcripts enriched in cell cycle processes (p=1.97e-83; Figure 3A) and other processes related to cell division (Supplementary Table S13) which closely track control and only prematurely decrease at the end of the drought treatment at 29 DAS before tracking control at rewatering. No overshoot is detectable (Figure 3A). Drought responsive transcripts behave atypically in clusters 2 and 3. Transcripts in cluster 3 enriched in cell wall organization and biogenesis (p=4.4e-06) increase in drought with a peak at 29 DAS and drop upon rewatering and they decrease with age in control conditions (Figure 3A). Transcripts in cluster 2 increase beyond the drought treatment into the first time point after rewatering before dropping to levels comparable with control and are enriched in RNA modification (p=4.1e-79). Both clusters 2 and 3 show opposite behaviour during progressive drought and progressive aging. Different clusters deviate from the trajectory of increasing or decreasing control abundances at different times. While cluster 4 increases early, cluster 1 deviates on 25 DAS and cluster 5 on 27 DAS. The decreasing clusters 6 and 7 deviate early while cluster 10 only deviates at 29 DAS (Figure 3A). Clusters 1, 4, 6, 7, 8, 10 track or inverse track the patterns of the ABA synthesis pathway and signaling although amplitudes are different (Figure 3A, Figure 2A). Clusters 4, and 9 track the anthocyanin pathway (Figure 3A, Figure 2B). Clusters 2, 3, and 8 represent different programs based on showing different patterns.

**Figure 3.**
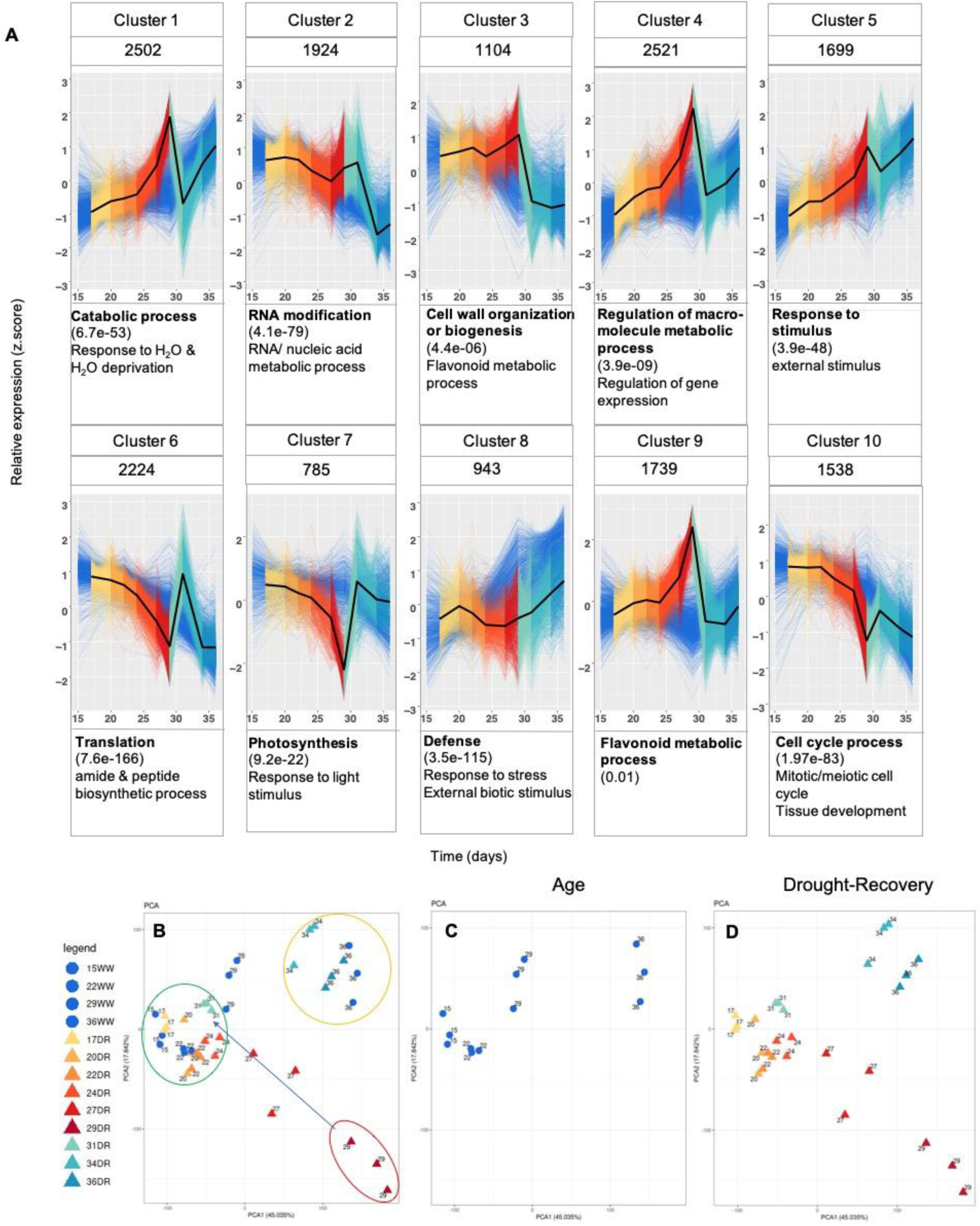
Pattern analysis of transcriptome data. **(A)** PAM clustering of 16,979 transcripts in ten clusters with drought time series in color gradient, its median in black, and control in dark blue; transcipts count atop, selected GO terms below, most significant GO term in bold with all GO terms available in Supplementary Tables S4-S13 (**B-D)** PCA of all non-zero transcripts with drought replicants (DR) are in triangles, well-watered (WW) controls as circles. Split view of PCA analysis showing (**C**) control samples only and **D)** drought and rewatering samples only.

To identify the dominant patterns in the complete transcriptome dataset, dimensionality was reduced to two dimensions via principal component analysis (PCA) (Figure 3B-D). The first two components represented 45% of the variation and 17% of the variation, respectively. In the first principal component data points separate based on plant age in control conditions and in drought conditions until 29 DAS (Figure 3B). In control conditions, the replicates of all 4 samples group with small separation between 15 DAS and 22 DAS and progressively larger separation to 29 DAS and 36 DAS (Figure 3B, C). Under drought treatment, premature aging is obvious with 29 DAS exceeding 36 DAS in control conditions and 27 DAS exceeding 29 DAS in control conditions (Figure 3B, D). Samples 27 DAS and 29 DAS also separate in the second principal component (Figure 3B, D) in opposite directions as aging control samples, a likely consequence of the patterns observed in cluster 2 and 3 where transcripts behave disparately during drought and aging (Figure 3 A). Upon rewatering, drought treated samples “jump back” towards young control samples before almost catching up with 36 DAS control samples at 36 DAS of drought and rewatering treatments (Figure 3B). The cluster and PCA analysis demonstrate that the patterns shown for the ABA synthesis and signaling transcripts re-occur in other transcripts which represent the majority of variation in the dataset. They also show that the programs underlying the aging of the rosette and the drought response overlap to a large degree (Figure 3).

### Majority of DEGs peak at the end of progressive drought stress

To quantify the transcript changes during drought treatment and rewatering compared to young, well-watered plants, differentially abundant transcripts were determined with edgeR resulting in 12.710 upregulated transcripts (log2FC > 0; FDR < 0.01) and 11.873 downregulated transcripts (log2FC < 0; FDR < 0.01) of 27,416 transcript analyzed (Figure 4, Supplementary Table curated_mothertable) An Upset plot showing the twenty intersections with the largest number of differential transcripts out of 512 possible combinations was generated to further dissect the shared and singular differential transcript expression for each day during the progressive drought stress and recovery (Figure 4). The combined abiotic and internal signals of drought stress and aging resulted in progressively more differentially abundant transcripts peaking at the last day of drought stress with about 12000 deregulated genes of which 2178 were specific to that time point only (Figure 4). 503 additional genes are specific only to the late time points 27 and 29 DAS of the drought stress treatment (Figure 4). GO term enrichment analysis of these 2.681 genes show significant GO terms for ‘metabolic process’ (1e-17), ‘regulation of gene expression’ (7.15e-15), ‘response to water‘ (1.05e-07) for 2324 upregulated genes. For 357 downregulated genes, ‘response to light stimulus’ (5.2e-17), ‘response to abiotic stimulus’ (2.69e-10) and ‘photosynthesis’ (4.03e-10) are among the top 10 GO terms (Supplementary Figure 3). Upregulated genes at the last day of the progressive drought stress showed the following unique GO terms: ‘nitrogen compound metabolic process’, ‘regulation of gene expression’, ‘cellular response to ABA stimulus’, ‘shoot system development’, ‘flavonoid metabolic process’. Whereas GO terms ‘photosynthesis’, ‘response to cold’, ‘chlorophyll metabolic process’, ‘mitotic nuclear division’ and ‘response to salicylic acid’ were observed unique to downregulated genes for drought (Supplementary Figure 4B). To analyze the changes with the largest amplitude at 29 DAS and 14 days of progressive drought treatment, the top 50 of upregulated genes (Table 2) and downregulated genes (Table 3) were investigated. Fifteen genes are involved in signaling (e.g. response to ABA, water, water deprivation, auxin), eight genes are involved in transcriptional regulation, seven genes are involved life cycle processes (seed/root development, leaf senescence), four genes are associated with carbohydrate metabolic process and protein modification. Three genes are involved in defense, transport and biological processes each. And lastly, represented with one gene in each category, plant VAMP (vesicle-associated membrane protein) family protein (cellular organization, protein of unknown function, DUF599 (secondary metabolic processes) and 3-ketoacyl-CoA synthase 2 (fatty acid biosynthesis process) are among the top 50 upregulated genes on day 29 but downregulated on day 31.

**Figure 4.**
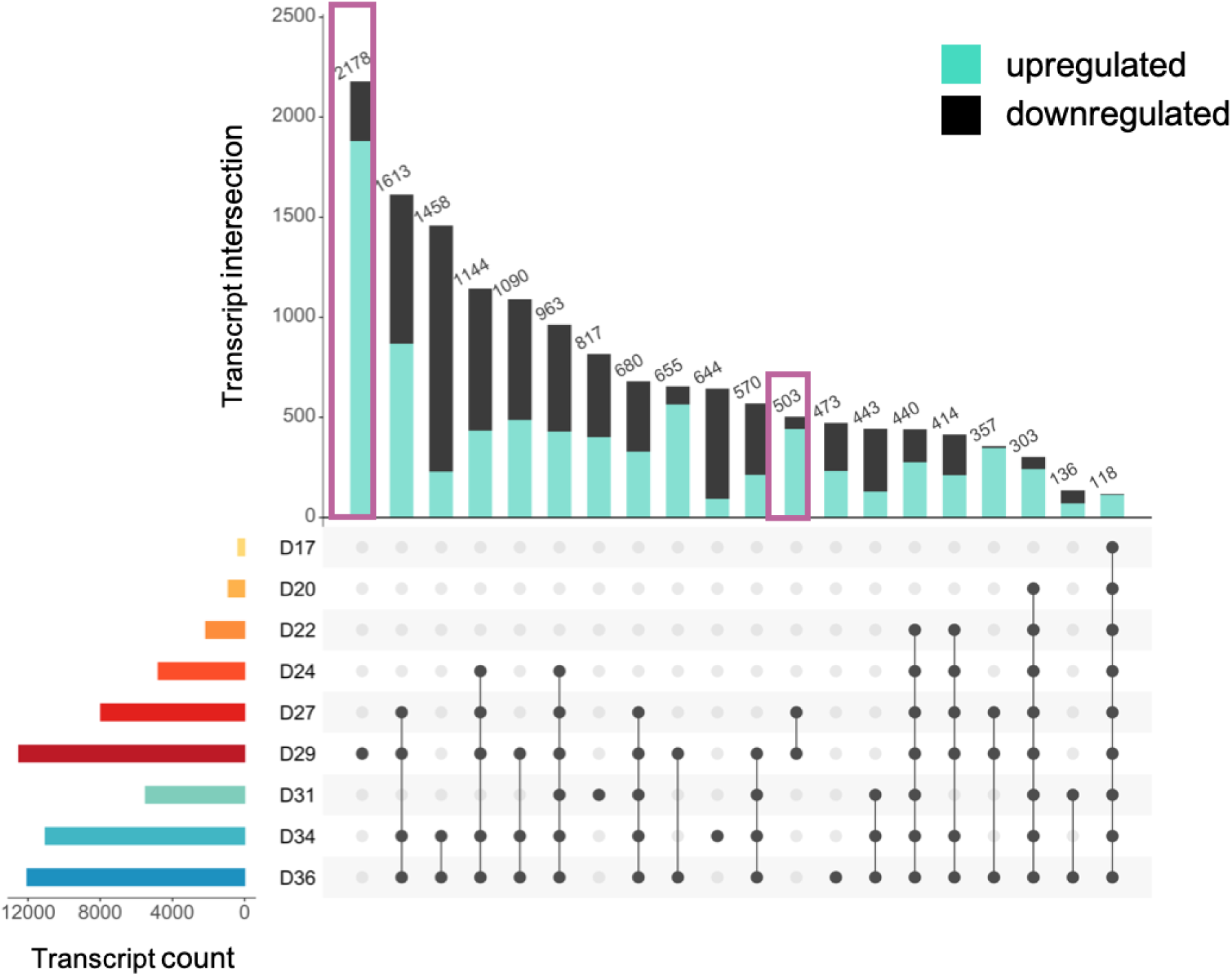
UpSet plot of differentially regulated transcripts across the progressive drought/rewatering treatment. Left bar plot labeled “Transcript size” displays the total number of differential transcripts compared to 15 DAS; The occurrence of transcripts within intersection as lollipop plots are represented in the top barplot, in which the upregulated transcripts were displayed in dark teal.

The fifty genes downregulated at the end of the progressive drought treatment include eleven genes involved in signaling. Here ‘response to light stimulus’, ‘cold’ and ‘gibberellic acid mediated pathway’ are downregulated at the end of the progressive drought (day 29). Six genes are involved in photosynthesis and life cycle respectively. The latter contains the transcription factor ‘response regulator 4’ (AT1G10470) that is involved in the regulation of the circadian rhythm. Four genes are part of defense, whereas 3 belong to protein modification process and carbohydrate modification. The latter contains genes relevant for cell-wall modification such as ‘expansin A8’ (AT2G40610) and Pectin lyase-like superfamily protein (AT5G63180). Two genes each can be associated with transcription regulation fatty acid biosynthesis, transport, nitrogen biosynthesis and transcription regulation. The latter contains two transcription factors ‘BR enhanced expression 2’ (AT4G36540) and ‘bZIP transcription factor family protein’ (AT5G28770.2). One gene can be associated with the pigment metabolic process, secondary metabolic process, amino acid modification and biological process. Surprisingly, transcripts frequently associated with drought stress such as those involved in reactive oxygen species detoxification (ROS) and late embryo abundant proteins (LEA) hypothesized to protect during desiccation are absent from these drought specific top50 lists.

### LEA and ROS show a diverse response towards progressive drought stress

A total of 246 ROS-associated transcripts (Oliveira et al., 2019) were analyzed in detail. Out of 10, 9 superoxide dismutases (SODs) passed the “differential transcript abundance in at least one contrast filter, of which 2 showed downregulation during the entire progressive drought stress (CCS, CSD1), 3 show upregulation during the drought stress with exception of day 27 and 29 (CPN20, FSD3, FSD2). CSD2 shows a turning point at day 22 from upregulation to downregulation. The remaining 3 displayed mostly upregulation throughout the progressive drought stress (Supplementary Table curated_mothertable). Transcripts for MSD1 show the highest abundance at day 20, for FSD1 at day 27, and only for a single SOD, CSD3, at day 29 (Supplementary Figure 5). Four genes belonging to ascorbate peroxidases (APX) are downregulated at the last day of progressive drought stress (APX4, APXT, APXS, APX5). The latter shows its highest expression on day 27. For 3 APXs (APX1, APX3, APX6) increased abundance is observed at day 29 of which APX6 shows the highest expression among the family. The expression patterns of MDAR2 and MDAR4 are almost identical with downregulation for the majority of the stress treatment followed by upregulation at the last two days of the stress time series of which day 29 shows the highest expression respectively. MDAR6 and MDHR show a reverse expression pattern from the previous two, here increased abundance is observed until day 22 and 24 respectively followed by a downregulation on days 27 and 29 (MDAR6) or on days 24 up to 29 (MDHR) (Supplementary Table curated_mothertable). With 37 members, the peroxiredoxins are the biggest family belonging to ROS-related enzymes. Fifteen enzyme-coding transcripts show upregulation towards the end of the progressive drought stress (PER11, PER15, PER17, PER20, PER22, PER25, PER29, PER3, PER34, PER34, PER42, PER49, PER51, PER52, PER53, PER64, PER67, Supplementary Figure 6). For 14 enzyme-coding transcripts we observe upregulation at the beginning of the progressive drought stress followed by a downregulation at the end (PER12, PER16, PER19, PER23, PER31, PER32, PER35, PER37, PER4, PER47, PER63, PER65, PER72, PER9). Five show downregulation throughout progressive drought stress treatment (PER21, PER33, PER39, PER50, PER71) (Supplementary Table curated_mothertable). For 27 enzymes belonging to GR the majority of transcripts shows downregulation towards the end of the progressive drought stress (GRC11, GRXC4, GRXC5, ROXY1, ROXY2, GRXS1, GRXS10, GRS11, CXIP1, CXIP2, GRXS2, GRXS3, GRXS4, GRXS5, GRXS6, GRXS7, GRXS8) whereas 8 display an upregulation at day 29 (GR, GRXC1, GRXC10, GRXC13, GRXC14, GRXC6, GRXS17, GRXS9). GRX480 displays its lowest abundance at day 24 whereas GRX4 shows its highest abundance at day 20 (Supplementary Table curated_mothertable). All enzymes belonging to AOX and CAT protein families show transcript upregulation at the end of the progressive drought stress (day 27 and or day 29). For AOX1C and HAOX2 the highest abundance is displayed at the onset of the recovery phase (day 31), whereas for the remaining members we see an opposite effect when rewatering is introduced to the plants (Supplementary Figure 7) (Supplementary Table curated_mothertable). In summary, while transcripts of specific isoforms of ROS detoxifying enzymes are indeed upregulated at some point during drought stress, a general upregulation of ROS detoxifying enzymes cannot be detected in the data neither at the initial more moderate nor at the later severe, but still sublethal drought stress.

For the majority of transcripts encoding the 51 LEA proteins in Arabidopsis (Hundertmark and Hincha, 2008) an upregulation towards the end of the progressive drought stress (day 29) followed by an opposite downregulation with the onset of rewatering (day 31) can be detected (Supplementary Figure 8).

### The drought responsive TF network extends beyond ABA and anthocyanin

To test whether the transcript abundance patterns detected in the clusters 2 and 3 which represented patterns different from the ABA and anthocyanin pathway patterns are associated with any transcription factors known to associate with drought stress independent of ABA, we examined their expression patterns (Figure 5). Predominant representatives within the ABA-dependent pathway are: NF-Ys, MYBs, AITRs and WRKYs. The NF-Ys TFs (NF-YA3, NY-YA5, NF-YA7, NF-YA10, NF-YB2, NF-C2) show uniform upregulation as drought stress treatment progresses. With the exception of NF-YA7 and NF-YC2 all show a dip in expression and a near catching up with the control plants after rewatering was initiated. The R2R3-MYB TFs MYB2, MYB94 and MYB96 show upregulation as drought stress advances.

**Figure 5.**
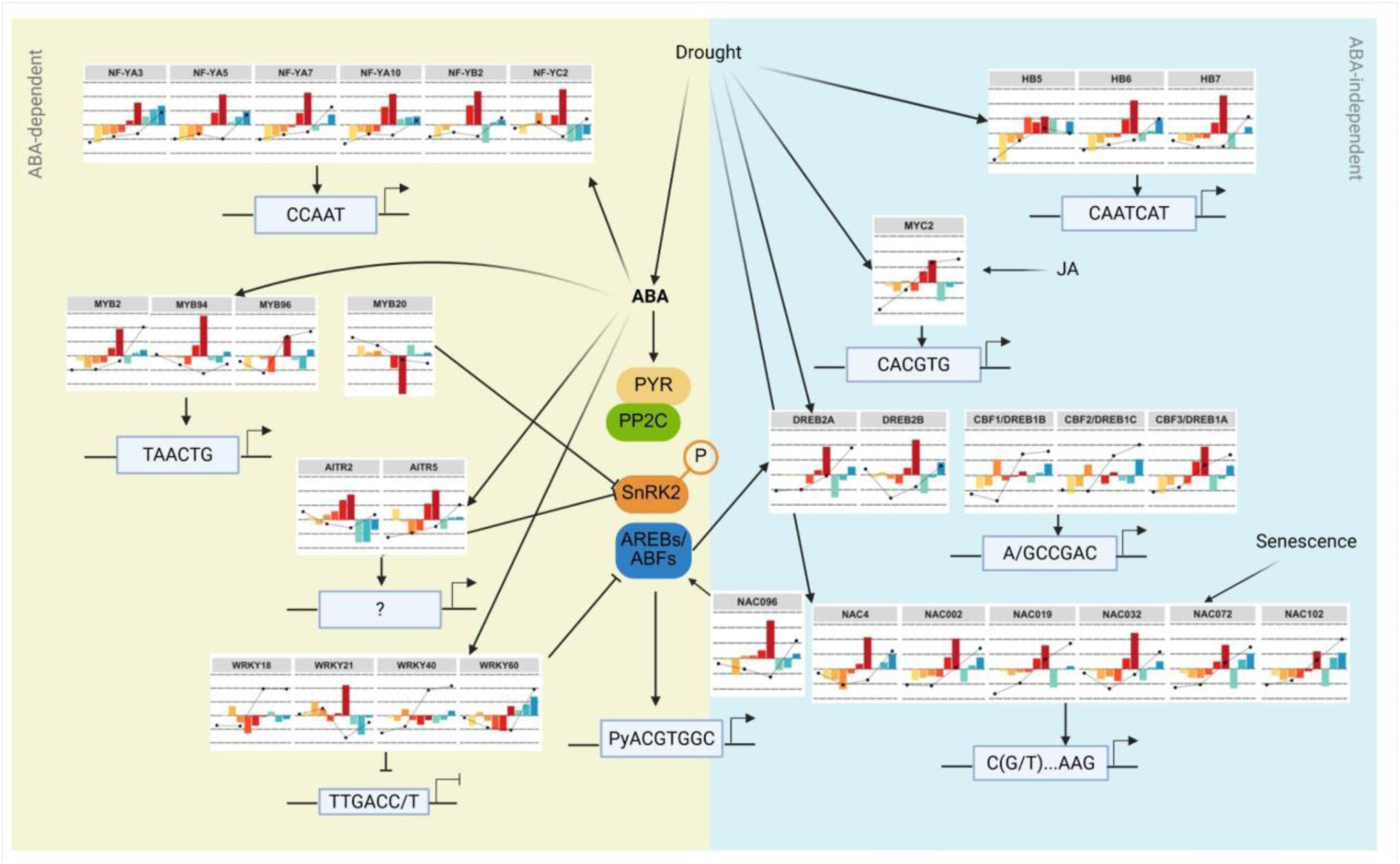
Model overview of TFs divided between ABA-dependent and independent signaling according to (Yao et al., 2021). The progressive drought stress is color coded from yellow-orange-red color scheme bars (one color per day), recovery is colored in green-blue bars. ABA-dependent signaling pathway is depicted in a yellow background whereas the ABA-independent signaling components are shown in a light blue background. Inhibitory regulations (other TFs or CREs) are indicated by a cross-bar, whereas positive regulations are depicted with an arrow. CRE-motifs are displayed in boxes (according to (Akhtar et al., 2012; Wang et al., 2021; Xie et al., 2010; Yao et al., 2021; Zenker et al., 2024). Figure was generated with BioRender.

For MYB20, reported to be a suppressor of PP2C (ABI2), transcripts show a downregulation towards the end of the progressive drought stress treatment (Cui et al., 2013). A novel TF, namely ABA-induced transcriptional repressors (AITRs), represses PP2Cs such as ABI2 upon drought stress. It has been reported that a double mutant of *aitr2 aitr5* shows strong drought tolerance (Tian et al., 2017; Yao et al., 2021). Here, both AITR2 and AITR5 transcripts show upregulation towards the end of the progressive drought stress followed by mirroring the expression upon rewatering. However, they do not catch up in expression as plant ages (Figure 5). The WRKY-type TFs WRKY 18 and 40, are known to negatively regulate AREB/ABFs, show overall a downregulation towards the end of the progressive drought stress compared to the ones previously observed (Figure 5, Supplementary Figure 9) (Yao et al., 2021). However, for WRKY 21, 28 and 60 upregulation on 29 DAS was observed (Figure 5, Supplementary Figure 9).

Within the ABA-independent signaling perception of drought stress, the bHLH TF MYC2 and the NAC-type TF NAC096 show similar behavior compared to HB6 and 7 and the ABA biosynthesis and signaling transcripts (Figure 5, Figure 2A). Of the HD-leucine zipper family TFS HB5, 6, and 7, HB6 and HB7 are progressively upregulated with drought, drop below baseline upon rewatering, and catch up with controls at 36 DAS (Figure 5). HB5 increases with drought stress without rebounding upon rewatering (Figure 5, Supplementary Figure 9). In contrast, the ERF/AP2 TFs of the CBF/DREB families show diverse patterns under drought but not during aging during which they all increase. CBF3/DREB1A and DREB2A have the ABA-type pattern under drought and rewatering, while CBF1 and CBF2 have diverse patterns unrelated to the ABA-type pattern or cluster 2 or 3 patterns (Figure 5). The senescence associated NAC-type TFs (NAC4, NAC002, NAC019, NAC031, NAC072, NAC102), which have been associated with drought, have a similar pattern compared to the ABA-type under control conditions although two of them dip before rising with age (Figure 5). All peak at 29 DAS, dip initially during rewatering, and catch up (Figure 5, Supplementary Figure 9).

### Phenotypic stages of drought stress manifestation are associated with largely disjunct sets of genes

In order to identify transcriptomic markers of the observed phenotypic changes with different time of onset in response to drought stress, we fitted a random forest regression to predict the course of each of the 20 relevant phenotypic traits identified above given the course of all 17k differentially expressed genes and again used the Boruta algorithm to select genes with strong predictive power. In total, 864 genes were found to be predictive, 428 of which were associated with architectural traits, 297 with FLUO color traits, 187 with RGB/VIS color traits, and 25 with the NIR emission trait (Supplementary Table S14). Phenotypic traits were highly predictable based on the expression of selected genes with a median cross-validation R^2^ value of 0.86 (Supplementary Table S15). The connections between genes and traits were visualized in a network using a force-based layout algorithm (Figure 7A). The different trait categories formed distinct subnetworks with only 67 of the 864 genes being predictive for traits from more than one category. The fluorescence subnetwork is very dense, indicating many shared predictive genes among the traits, while traits in the architectural subnetwork are more loosely connected. The NIR trait is centrally positioned, forming connections with traits mainly from the architectural and fluorescence categories (Figure 7A).

To gain an insight into the functional significance of the predictive transcripts enrichment of Biological Process GO Terms was evaluated. For the upregulated genes within the overall prediction network, 212 GO terms were significantly enriched (Supplementary Table S16) with the most strongly enriched terms belonging to the response to stimulus category (GO:0050896); specific terms included response to stress (GO:0006950), defense response (GO:0006952), and response to oxygen-containing compound (GO:1901700). Other terms were related to coloration processes: pigment metabolic process (GO:0042440), flavonoid metabolic process (GO:0009812), and anthocyanin-containing compound metabolism (GO:0046283). The architectural subnetwork was mostly enriched with stimulus and defense response terms while the fluorescence subnetwork had stronger enrichment in the pigmentation and color-related terms than the complete network. It also contained the term leaf senescence (GO:0010150). The VIS category was most strongly enriched in terms regarding starvation response (GO:0042594) and nutrient levels (GO:0031669) as well as cellular detoxification processes (GO:1990748). The top hits in the NIR category were lignin processes (GO:0009808) and iron ion transport (GO:0006826). The downregulated genes of the entire prediction network were also most strongly enriched response terms for biotic stress factors (such as response to other organisms, GO:0051707 or defense response, GO:0006952). They also contained developmental terms such as regulation of cell cycle (GO:0051726) and leaf development (GO:0048366). These defense and immune response terms were also significantly enriched within the architectural, VIS and fluorescence color subnetworks and the latter was also enriched with growth and tissue development terms like trichome morphogenesis (GO:0010090) and plant epidermis development (GO:0090558) as well as cell cycle terms (regulation of cell cycle process (GO:0010564) and mitotic cell cycle phase transition (GO:0044772).

Given the strong separation of subnetworks we hypothesized that genes within each subnetwork follow a similar expression pattern, each corresponding to a distinct phase in the phenotypic manifestation of drought stress response. However, genes within a subnetwork do not seem to follow a common pattern and all subnetworks contain genes following early, late, and overall different expression patterns (Figure 7B) although there are significant enrichments in genes belonging to certain PAM clusters (Figure 3A) in the different subnetworks (p-values < 0.0001, Figure 7C): While the architectural traits are largely associated with genes belonging to clusters 2, 5, 6, and 8, the genes associated with fluorescence traits stem mostly from clusters 1, 2, 4, and 10. The traits in the VIS subnetwork are significantly enriched with genes from clusters 7 and 9 and the 25 transcripts associated with the NIR emission trait belong to many different PAM clusters.

Given the strong predictive power of the selected genes for the emergence of their associated phenotypic traits and the enrichment in relevant biological processes, we tested if the algorithm selected causal candidate genes or biomarkers. We therefore ranked genes using multiple criteria (number of associated traits and subnetworks, strength of association) to cover different scenarios and filtered the top 230 ranked genes for the availability of two independent insertions from the SALK and GABI-Kat collections (Supplementary Table S2). We propagated 71 of these knockout lines together with the Col0 wildtype and confirmed the expected insertion to be present in homozygous state using PCR (Supplementary Table S3). This yielded 22 knockout mutants targeting 11 genes (2 independent insertion events per gene) which were subjected to the same experimental conditions and measurement protocols alongside the background wildtype (Supplementary Table S17). While some knockouts showed small differences in their drought stress phenotype compared to the wildtype, none of the 11 genes tested in two independent replicates produced a significantly different phenotype of the predicted traits in both corresponding knockouts (Supplementary Figure 9). This indicates that our gene-to-phene approach did successfully identify biomarkers but did not select for causal genes.

## Discussion

### Drought induces distinct phenotypic and transcriptional programs

To analyze the programs underlying mild, medium, and strong, but nonlethal drought stress we employed a combined phenotyping and transcriptome approach with daily phenotyping, and three RNA-Seq samples per week (Figure 1A). This dense sampling allows a detailed, pattern based analysis for phenotypes and differential gene expression. The random-forest based Boruta feature selection algorithm identified 20 phenotypic traits predictive for drought stress in *Arabidopsis thaliana* (Figure 1C). We show that the drought stress response manifests in separable stages corresponding to different types of phenotypic traits: changes in the NIR intensity trait first occur in the early phases of drought stress, days before any differences are visible (Figure 1C, D) and with only very limited changes in the transcriptome (Figure 4). This NIR intensity trait is dependent on plant water content (Berger et al., 2010; Knipling, 1970), and has previously been demonstrated to be an effective monitor of drought stress levels (Chen et al., 2015). As the stress progresses, differences in color become apparent in both the visible and then fluorescence light spectrum (Figure 1C, E, F), likely due to changes in plant pigmentation. The plant might produce specific pigments for their antioxidative properties that are helpful to limit the damage by reactive oxygen species (Dixon’ and Paiva, 1995; Nakabayashi et al., 2014). Anthocyanins are intrinsically fluorescent molecules (Chanoca et al., 2018) and the transcripts which encode the enzyme that produce them are detectable at 27 and 29 DAS (Figure 2B). In line with earlier studies on the phenotypic effects of drought stress, biomass and other architectural traits are affected last during the longest and most severe drought (Chen et al., 2015; Harb et al., 2010). The progression of phenotypic traits follows a sequence which may reflect the plant defenses against the stress. Initially, there is no growth limitation (Figure 1B), only very limited changes in the transcriptional program (Figure 4), and no changes in visible or fluorescence traits predictive for drought stressed plants (Figure 1C, E, F). With increasing drought stress duration and intensity, the plant reacts at the transcriptome level (Figure 4), possibly with post-transcriptional regulation (Fujita et al., 2009), and ultimately with altered phenotype at the visible and fluorescence level (Figure 1C, E, F). Architectural traits are not predictive for drought-stressed plants at this point (Figure 1B) even if the onset of changes with small magnitude begin to manifest (Figure 1G). Growth limitation is the last resort of the annual *A. thaliana*.

The onset of predictive traits at distinct time points raises the possibility of distinct transcriptional programs controlling the different traits which then may onset at a distinct time.

### Drought responses are controlled by at least four sets of transcriptional programs

The analysis of two known transcriptional programs, the ABA synthesis and signaling program as well as the anthocyanin biosynthesis program, reveals a distinct threshold onset only for the latter (Figure 2B). Among the anthocyanin biosynthesis genes at least one isoform demonstrates this threshold onset behavior (Figure 2B). Double mutants of *pal1* and *pal2* showed a deficiency in anthocyanin under high light conditions and thus imply that PAL1 and PAL2 have a redundant role in flavonoid biosynthesis (Huang et al., 2010), explaining why induction of one isoform is sufficient. All transcripts encoding biosynthesis enzymes decline after rewatering (Figure 2B), indicating that anthocyanin accumulation is not a phenotype associated with aging in the annual *A. thaliana*. The transcription factors known to control the pathway only partially mirror the patterns of the target genes. Some of the of MYB transcription factors, MYB11, MYB12, PAP1, PAP2, and the bHLH TT8 but not MYB111 display the same abrupt onset on 27 DAS, high abundance at 29 DAS, and rapid drop upon rewatering. In contrast, the obligate partner WD40 TTG1 and the alternative bHLHs GL3 and EGL3 have different patterns (Figure 2B). This indicates that during drought stress the WD40 TTG1 abundance is likely not a limiting factor in the complex which controls the induction of anthocyanin biosynthesis genes (Figure 2B). No induction of anthocyanin biosynthesis or regulators is detectable with increasing plant age in control conditions (Figure 2B). The production of anthocyanins can also be induced by high light (Dubos et al., 2008) where it acts as a light attenuator or antioxidant (Zheng et al., 2021). It is likely that the intracellular CO_2_ limitation caused by closed stomata leads to electron acceptor limitation due to limited Calvin Benson Bassham cycle activity, and causes excess energy stress during standard light conditions. This explains why the transcriptional anthocyanin program clusters with genes in unusual patterns compared to others during progressive drought (Figure 3A). It is likely driven by consequences of drought, namely acceptor limitation in the Calvin Benson Bassham cycle and not by drought responsive transcription factors per se.

In contrast to the anthocyanin program, the ABA program has a gradual onset starting on 17 DAS and increasing until 29 DAS, the final time point during drought (Figure 2A). This suggests that the ABA program does not have a threshold onset but increases with increasing severity and/or increasing length of drought. In our experimental setup, drought was induced by water withholding and steadily increased in intensity making it impossible to distinguish between length and severity as factors (Figure 1A). The only transcripts encoding ABA biosynthetic steps dissenting from the gradual increase are the BCHs and ABA2 (Figure 2A). BCH2 is steadily expressed with increasing plant age and its level likely suffices for ABA synthesis. As the BCH product zeaxanthin is involved in processes beyond ABA synthesis such as non-photochemical quenching and the xanthophyll cycle (Young, 2006), its production is required throughout the life of a green plant. ABA2 is a short chain reductase and even small amounts are sufficient to allow ABA synthesis to proceed (Cheng et al., 2002). This may explain why no induction of ABA2 was detectable (Figure 2A). ABA2 also does not show any induction under short-term drought stress (Lin et al., 2007). The ABF transcription factors of the bZIP family downstream of ABA signaling show gradual induction as do the PP2Cs which rapidly shut down ABA signaling upon loss of signal and the SnRK2s which transmit the ABA signal at the post-translational level via phosphorylation events (Figure 2A). These bZIP family transcription factors target bind ACGT as the core motif (Zenker et al., 2024) and ABI5 is known to target photosynthetic genes for repression (Zhu et al., 2020). The ABA responsive TFs specifically bind the ABRE, ACGTG (Hattori et al., 2002). The bZIP transcription factor HY5 activates transcription via the G-box CACGTG (Toledo-Ortiz et al., 2014). It is thus likely that the observed downregulation of photosynthetic genes (Figure 3A) is regulated at least in part via the action of bZIP transcription factors which act as repressors on G-boxes. G-boxes are wide-spread in the Arabidopsis genome and extend beyond photosynthetic genes (Ezer et al., 2017). ABI5 can also act as an activator of gene expression for stress-responsive protein such as LEAs (Finkelstein and Lynch, 2000). As ABI5 can act as both an activator and repressor, there is potential for the other ABA-responsive bZIP TFs to act similarly and thus cause both up-and downregulation among the transcripts (Figure 3A) which follow or inversely follow the ABA pattern (Figure 2A). The drought response is also mediated via the DRE/CRT motif A/GCCGAC (Joshi et al., 2016) which is bound by a subset of the ERF/AP2 family (Zenker et al., 2024) that includes the CBF/DREB genes (Joshi et al., 2016). The two known CBFs CBF1 and CBF2 mediating the cold response are not included among the genes which respond to increasing drought (Figure 5) but the known drought responsive ERF/AP2s CBF3 and DREB2A are (Figure 5). In addition to these, NAC TFs, MYB-related TFs and HD-Zip TFs previously reported to be associated with the drought response respond to drought in a pattern similar to that of the bZIP TFs (Figure 5, Figure 2). Since all NACs bind a similar motif, the gapped CTT……AAG motif which enriches downstream of the transcriptional start site (Voichek et al., 2024), some MYB-related TFs bind the same motif (Zenker et al., 2024), and many HD-Zip TFs bind the same motif (Zenker et al., 2024), it is not surprising that strong, but sublethal drought ultimately affects a third of all genes in the Arabidopsis genome (Figure 4). Since drought induces the expression of TFs of many different families, there is ample potential for regulation via AND and OR rules using the known drought responsive motifs ABRE likely bound by bZIP-type TF ABI5, ABF1 and others, DRE and CE1 likely bound by the ERF/AP2-type TFs of the DREB and CBF families, and CE3 possibly bound by NAC-type or CAMTA-type TFs, and motif III. All these transcription factors have a pattern comparable with that of the ABA responsive bZIP types (Figure 2A, Figure 5). These TFs are hence likely relevant for the transcriptional programs similar or inversely similar to the ABA program in clusters 1, 4, 5, 6, 7, and 10. With the exception of the flavonoid responsive TFs (Figure 2B), the TFs driving the patterns in cluster 3, and 9 are currently unknown. The unusual pattern in cluster 2 where the effect continues into the rewatering phase is also currently unknown. The comparison of onset of predictive phenotypes for drought (Figure 1) with transcriptional patterns (Figure 2, Figure 3A) show that the onset of traits at least to some degree correlates with onset of linked transcriptional patterns. Growth depends on translation for protein production (Wu, 2024), and on cell division and expansion (Vercruysse et al., 2020). Transcripts related to translation are affected before transcripts related to cell division (Figure 3A) which coincide with the observation of growth arrest (Figure 1B). Anabolic metabolic pathways such as photosynthesis and chlorophyll biosynthesis decline while color and fluorescence traits become relevant for prediction (Figure 1C, Figure 3A). All these patterns have a gradual onset rather than a threshold onset observed for the phenotypic traits. It is thus likely that, with the exception of the anthocyanin biosynthesis, the phenotypic traits, despite their threshold onset for predicting drought (Figure 1), are controlled by transcriptional programs with gradual onset (Figure 3). The stages of phenotypic response are predicted by mostly disjunct sets of genes (i.e. subnetworks in Figure 7) and while specific gene expression clusters are enriched in each subnetwork, all subnetworks contain genes following early, late, and overall different expression patterns. This is likely because the random-forest algorithm underlying the Boruta feature selection approach can combine information non-linearly and unlike Figure 7, which shows z-score-normalized expression patterns, the algorithm learns specific TPM thresholds which makes genes with similar expression patterns not simply interchangeable (Breiman, 2001). This is also indicated by the different GO terms enriched in the upregulated genes of each subnetwork. Among the downregulated genes, biotic stress response and cell cycle/tissue development were prominent. Plants actively downregulate their capacities for disease response when subjected to drought stress (Fujita et al., 2006), mediated by the transcriptional activator AtMYC2 which is induced by ABA and represses jasmonic acid and ethylene-mediated pathogen defense genes (Anderson et al., 2004). ABA also downregulates growth-related genes (Cutler et al., 2010) which ultimately results in a smaller phenotype (Berger et al., 2010), so finding these terms enriched in the downregulated genes is plausible in the context of drought stress.

### Predictive genes are not necessarily causal

Even though the selected genes were strongly predictive of the emergence of drought-relevant phenotypes (Figure 6A, Supplementary Table S15), knocking them out did not significantly affect the predicted phenotype in any of the 11 tested cases (Supplementary Figure 9). Given the small difference (and sometimes even contradiction) between mutants targeting the same gene (Supplementary Figure 10), this is unlikely a case of insufficient statistical power. Since the Boruta feature selection algorithm finds sets of genes that are used in conjunction to predict the emergence of phenotypes rather than individual, independent genes (Kursa and Rudnicki, 2010), it is possible that knocking out a single gene at a time is not sufficient for affecting the expected phenotype but that manipulating multiple genes together would produce the expected result. It is also possible that the tested (and likely a majority of selected) genes are affected by a factor upstream in the gene regulatory pathway that also affects the phenotype, i.e. they are simply correlative but not causative. To find causal genetic loci rather than associated variation, it will likely be critical to limit the number of transcripts used for prediction based on prior biological knowledge as was successful with limitation to transcription factors previously (Halpape et al., 2023) and to account for functional redundancy (e.g. by using the SKM CUT tool from (Bleker et al., 2024)).

**Figure 6.**
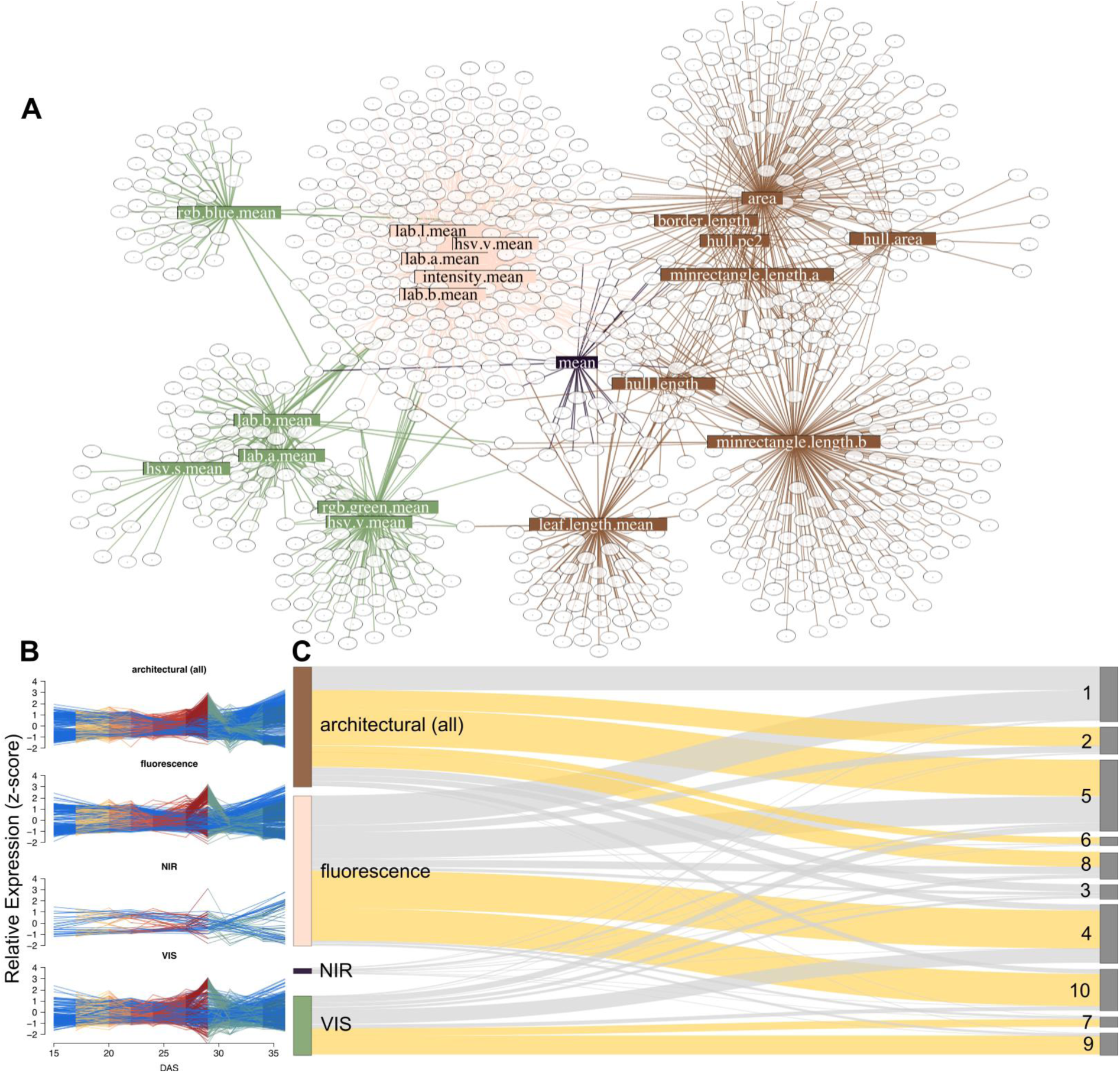
Transcripts associated with drought-responsive traits. **A)** Force-based network layout of relevant phenotypic traits and associated transcripts, colored by trait category. Traits of different categories clearly cluster together and form largely disjunct sub-networks. Gene names are omitted for visual clarity but are listed in Supplementary Table S14. **B)** Relative transcript abundance patterns of genes within each sub-network. **C)** Sankey diagram of genes in sub-networks and PAM expression clusters (Figure 3D). Dandelion colored connections symbolize significant enrichments of clusters in a subnetwork (all p-values < 0.0001, one-sided Fisher’s exact test, Benjamini and Yekutieli (2001) adjusted).

### Drought and maturation are coupled in the annual *A. thaliana*

Drought stress over a 14-day period shows progressive transcriptional changes in the second component of the PCA analysis (Figure 3B-D). With the onset of rewatering transcriptional changes “jump back” and abundances are similar to those from the beginning of the drought stress treatment (Figure 3B). This indicates that plants show premature aging at the end of the progressive drought stress treatment and rejuvenate to some degree with the onset of rewatering. The overlap in transcriptional programs is also obvious in the ABA program (Figure 2A) and seven out of ten clusters for significantly changed genes (Figure 3A). These effects may be due to single cells aging more quickly upon onset of drought, or the whole plant speeding up its lifecycle. Stress-induced flowering is a well-known (Takeno, 2016) and increasingly well understood (Katagiri et al., 2024) trait. The transcript abundance data indicates that an aging rosette in control conditions undergoes many of the same transcriptional changes as are brought on by drought (Figure 2A, Figure 3). Moreover, watering has a rejuvenating effect on at least 45% of the transcriptome (Figure 3B-D). Interference with drought response programs such as manipulating the expression of drought responsive transcription factors to increase drought tolerance may have unintended consequences on the speed of plant maturation and therefore yield.

## Data Availability

All data and code required for reproducing our results are freely available in our GitHub repository at https://github.com/Thyra/arabidopsis-boruta. Raw sequencing data were deposited in the National Center for Biotechnology Information (NCBI) and are publicly available via BioProject: PRJNA1155997. The raw RNASeq data are assigned to the following accessions: SRR30548061 -SRR30548099. Raw and processed imaging data are available at https://doi.ipk-gatersleben.de/DOI/f0a10c1e-ff92-48e4-a8a6-728b04a5b57b/ffba031d-eef2-49db-8380-b4fc46056d59/2/1847940088, including metadata compliant with MIAPPE v1.1 to ensure maximum transparency and reproducibility (Papoutsoglou et al., 2020).

## Supporting information

Manuscript Tables

## Author Contributions

EFC, DP, and AB wrote the manuscript. DP and AJ performed phenotyping experiments. EFC and DP performed data analysis and made figures. BW funded the RNASeq sequencing. All authors read and edited the manuscript.

