## Supplementary figures and images for "A sublethal drought and rewatering time course reveals intricate patterning of responses in the annual Arabidopsis thaliana"

### Supplementary Figure 1

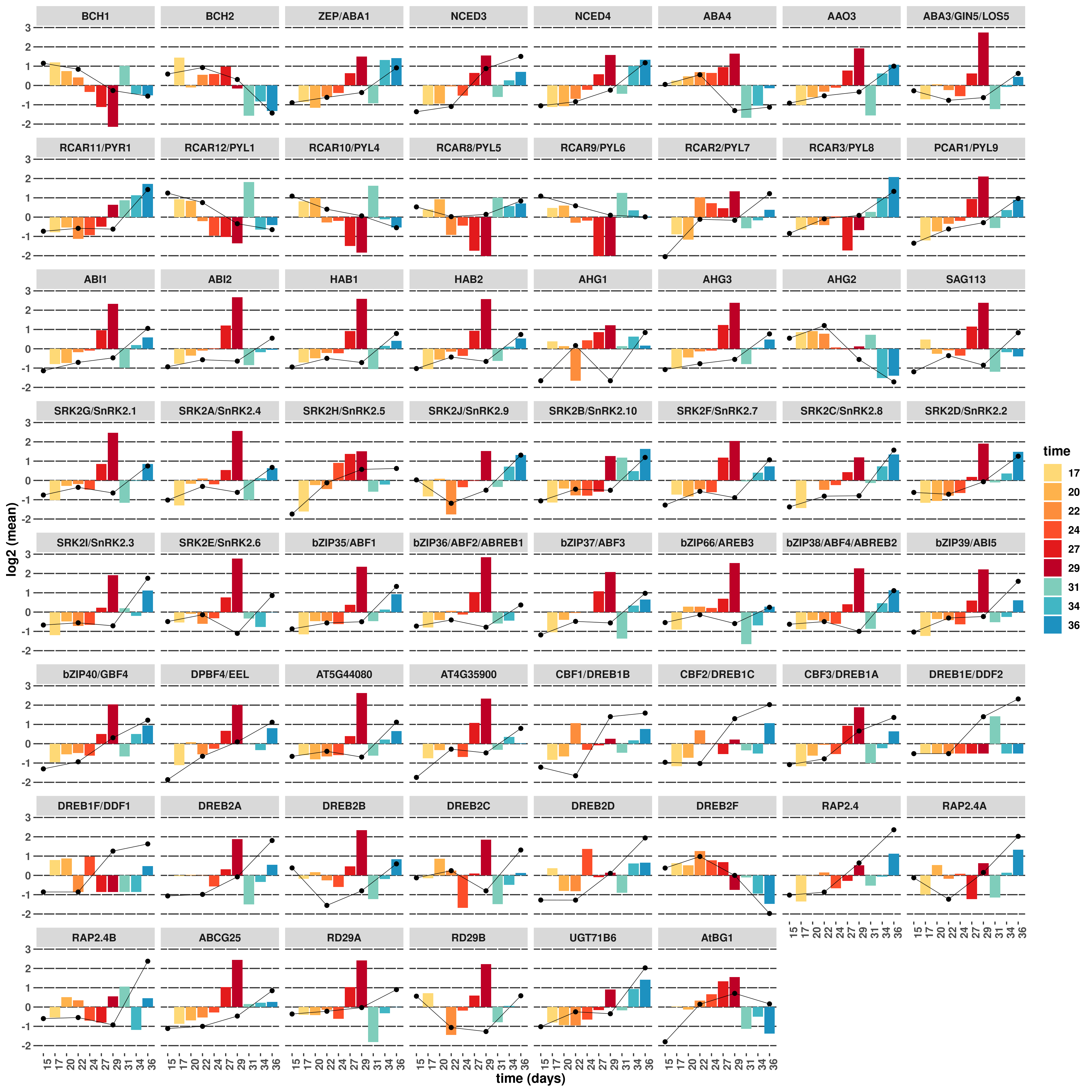

### Supplementary Figure 2

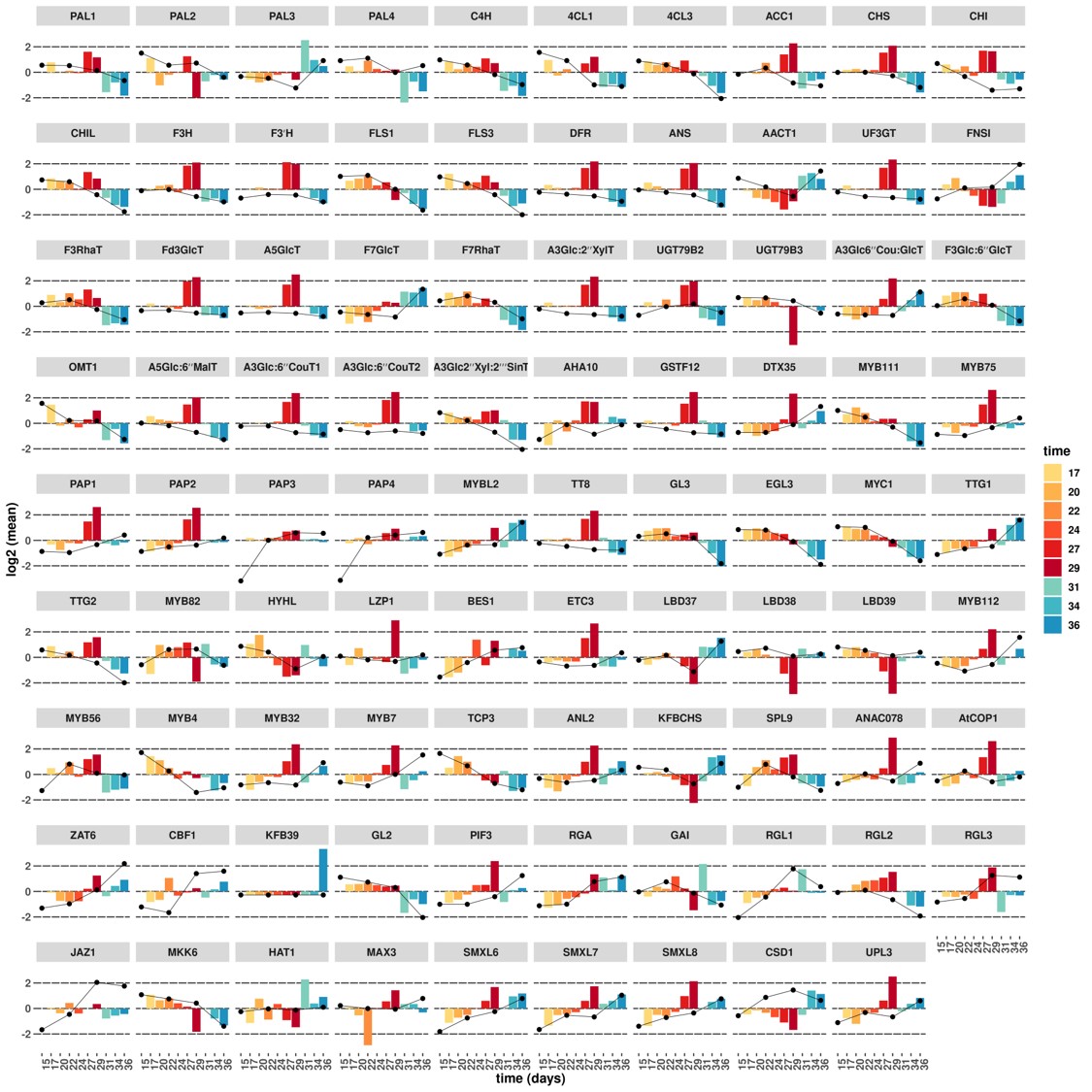

### Supplementary Figure 3

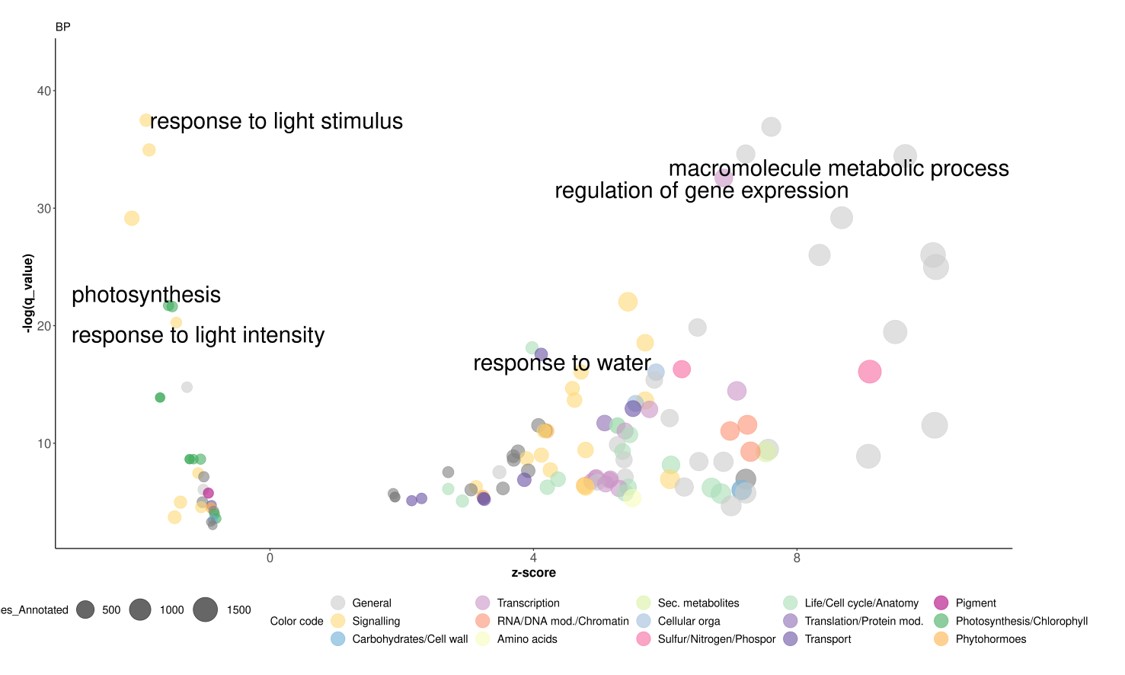

### Supplementary Figure 4

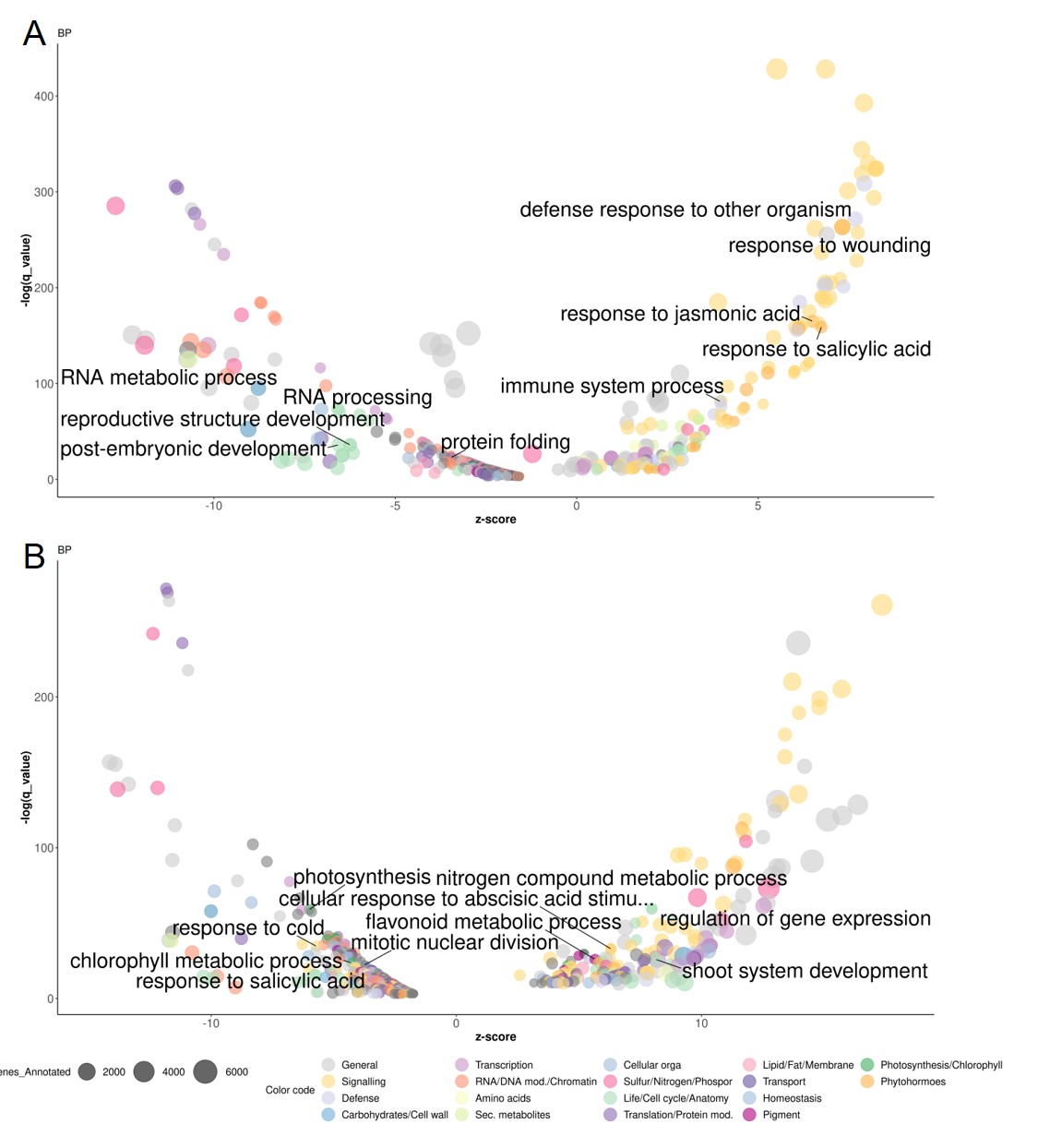

### Supplementary Figure 5

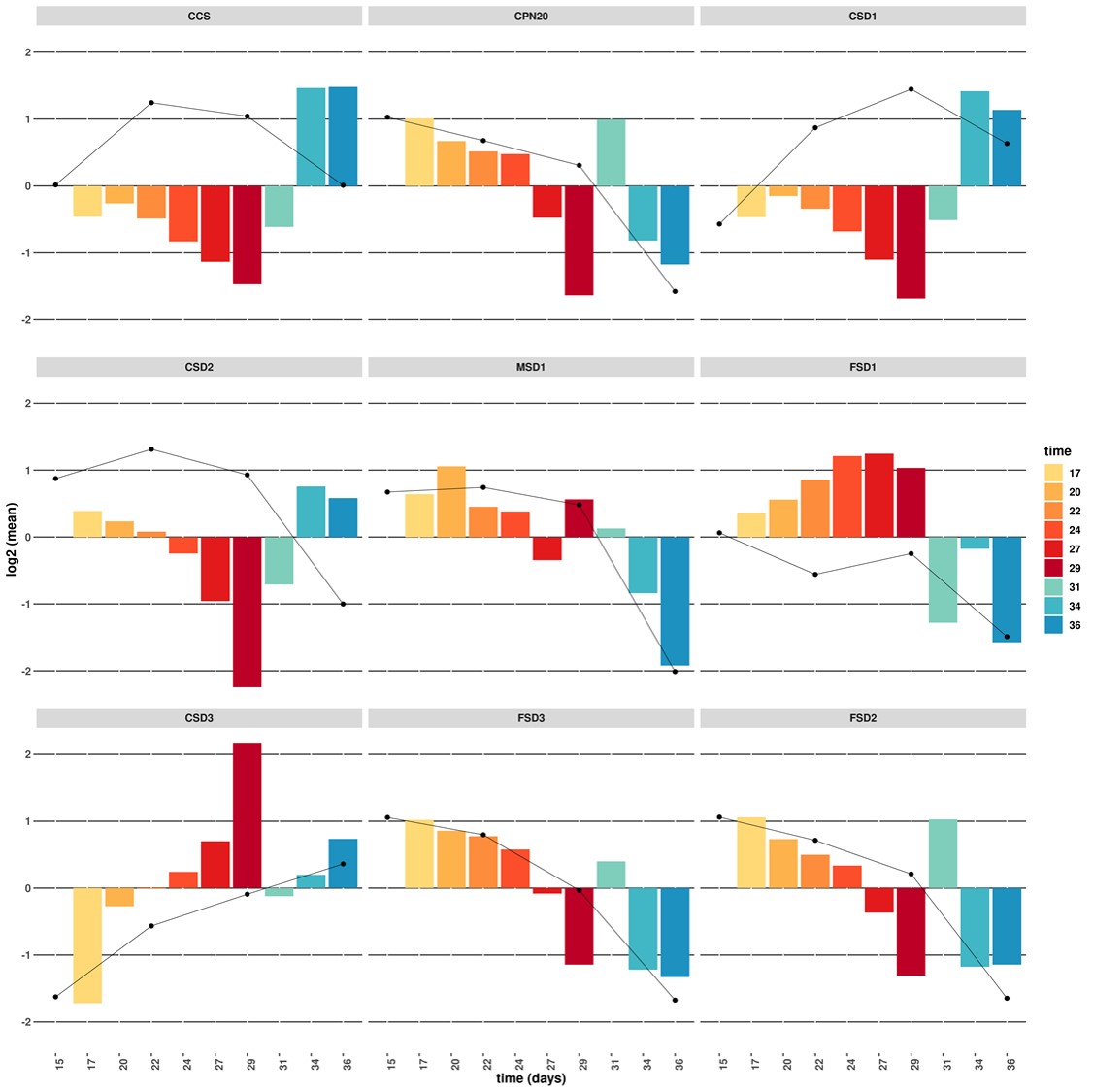

### Supplementary Figure 6

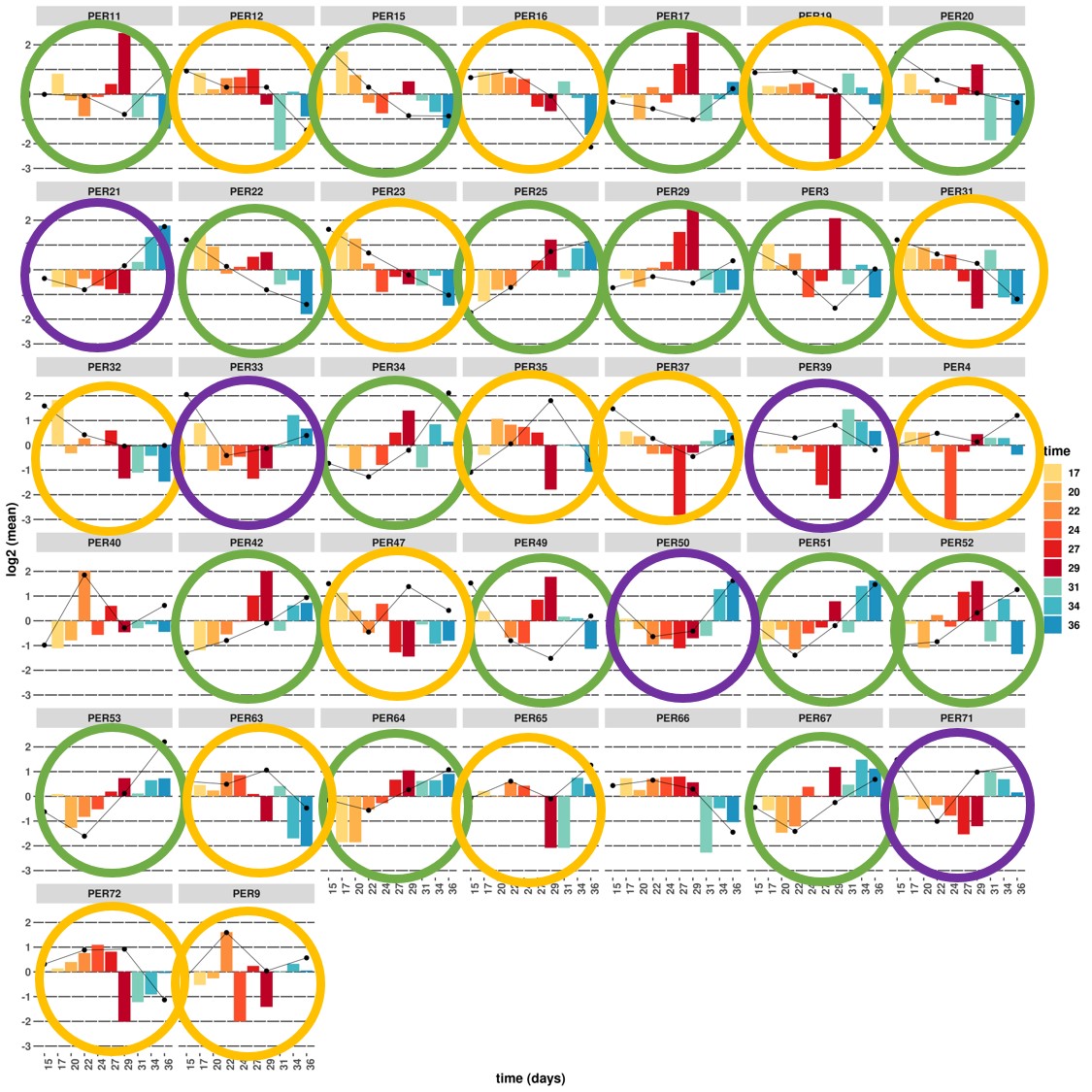

### Supplementary Figure 7

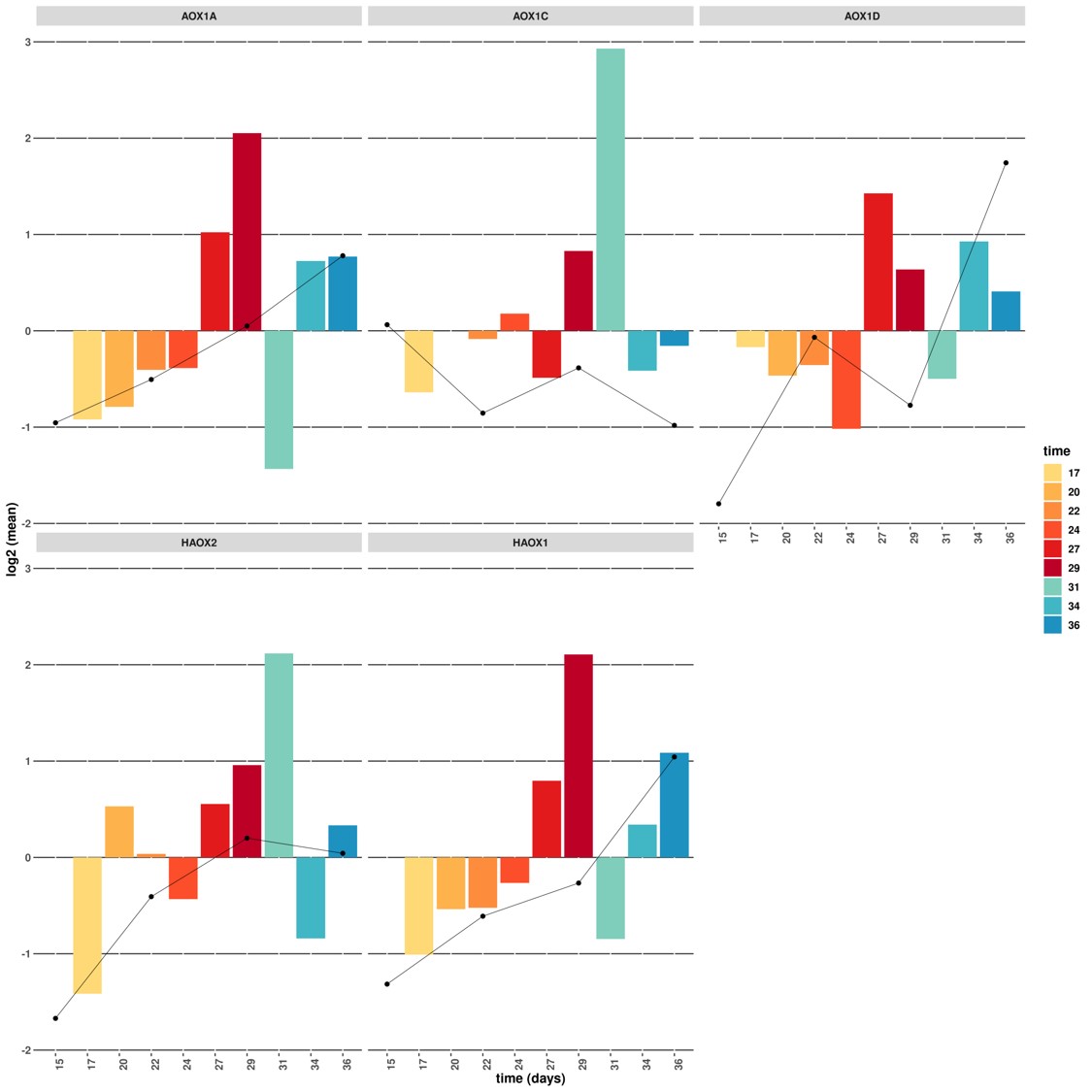

### Supplementary Figure 8

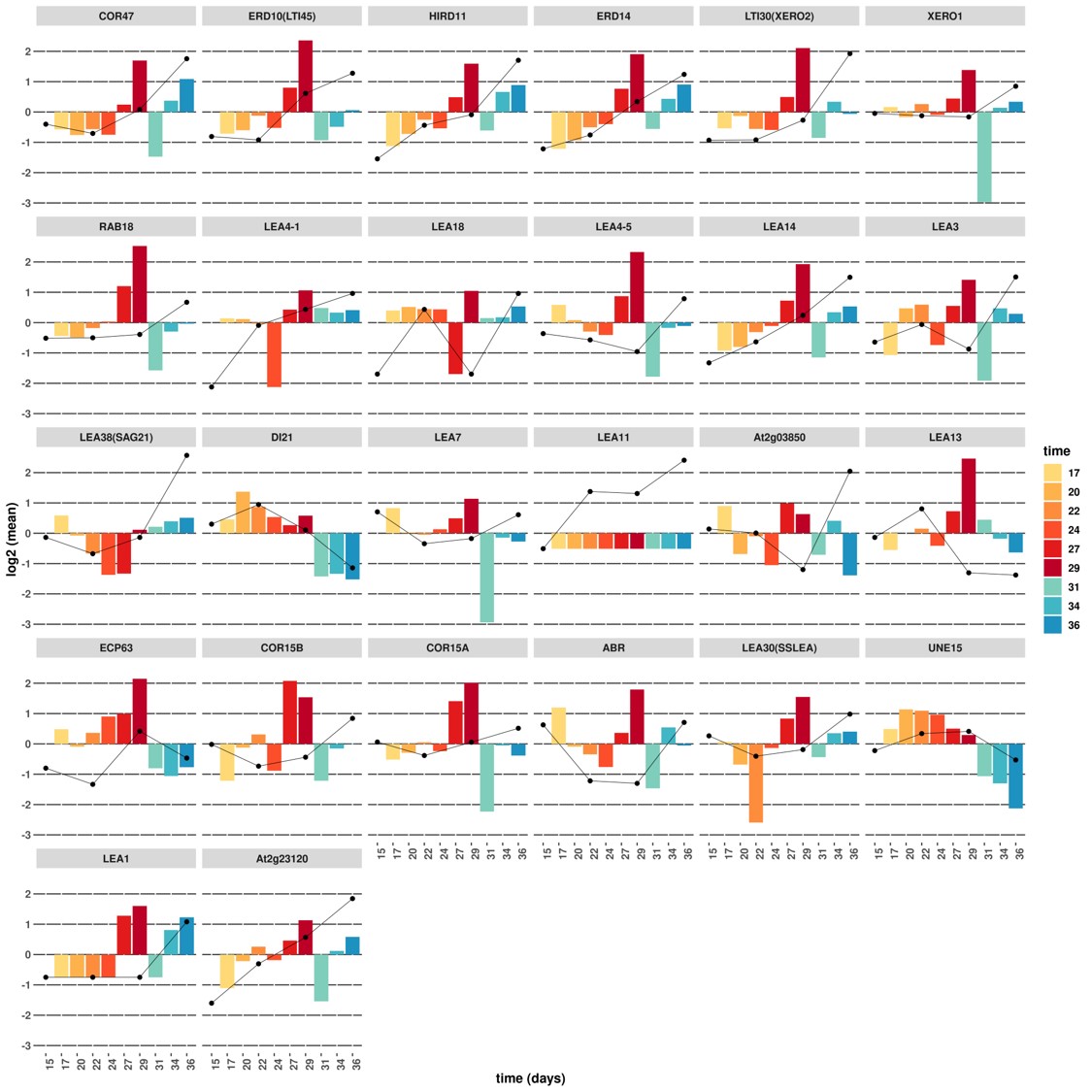

### Supplementary Figure 9

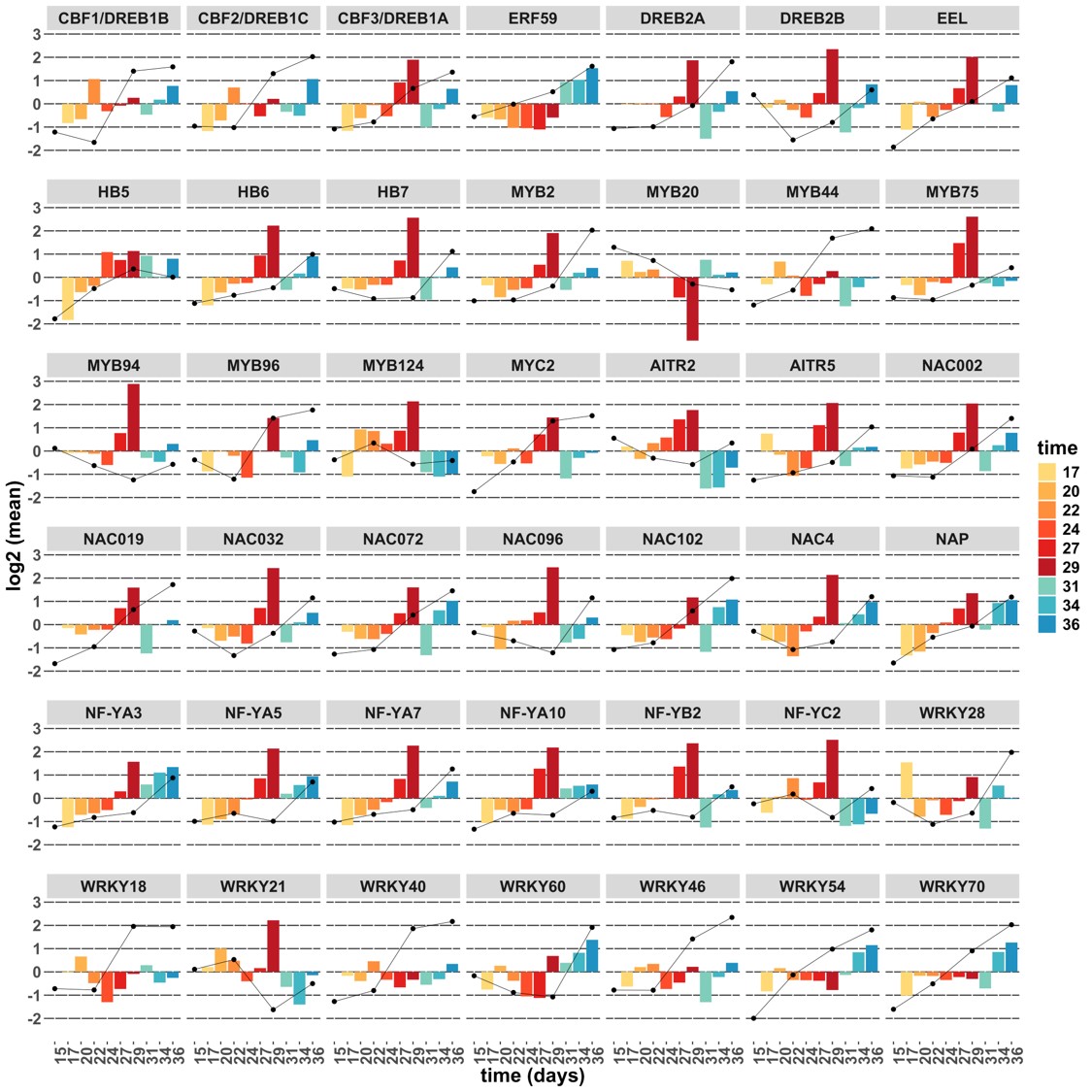
