## Supplementary Figure 10 for "A sublethal drought and rewatering time course reveals intricate patterning of responses in the annual Arabidopsis thaliana"

| Trait / Gene | AT3G51895 | AT5G03260 | AT3G05630 | AT5G54610 | AT3G06035 | AT3G45600 | AT3G16460 | AT2G39350 | AT2G44450 | AT2G18450 | AT4G11070 |
|---|---|---|---|---|---|---|---|---|---|---|---|
| top.geometry.fluo leaf.length.mean.px | | | | 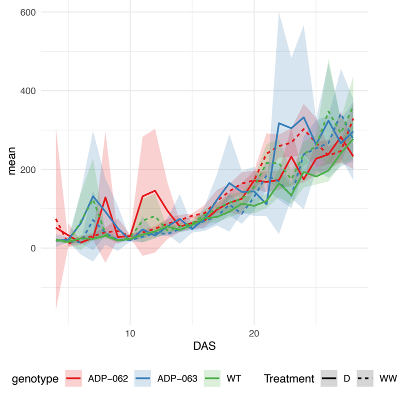 | | | | | | | |
| top.geometry.fluo minrectangle.length b.px | | | | 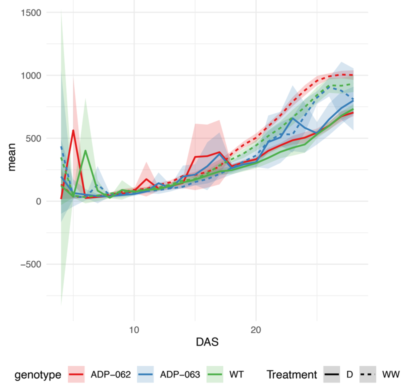 | | | | | | | |
| top.intensity.fluo hsv.v.mean | 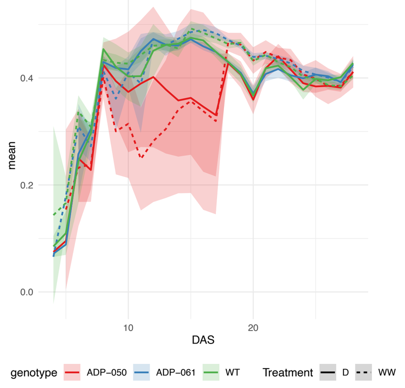 | | | | | 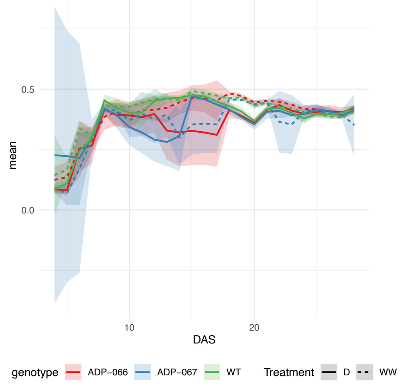 | | | 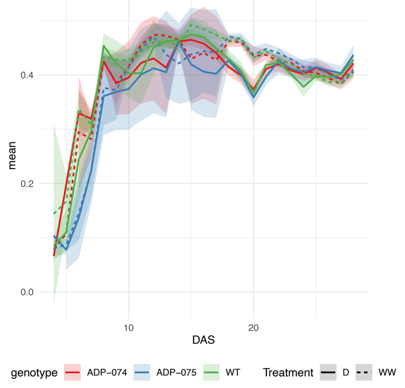 | | 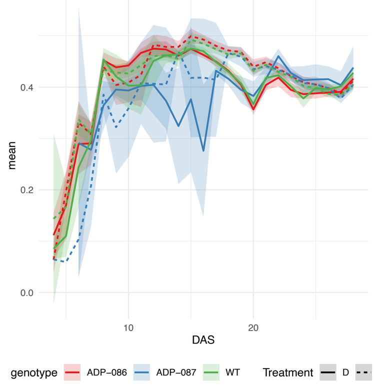 |
| top.intensity.fluo intensity.mean | 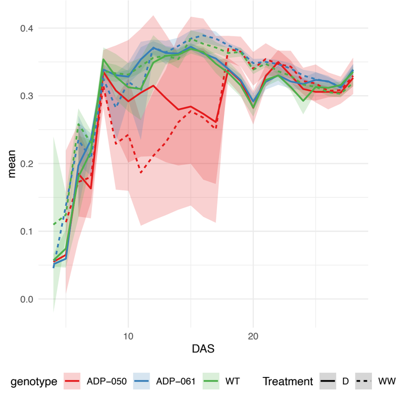 | | | | | 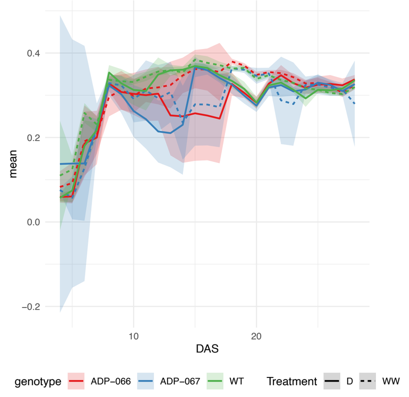 | | | | | 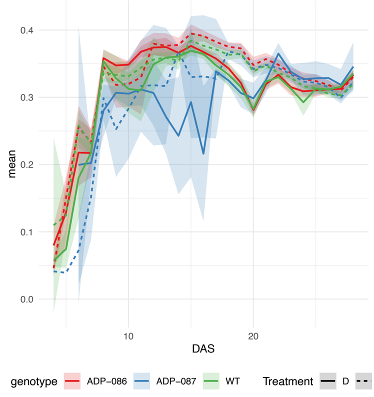 |
| top.intensity.fluo lab.a.mean | 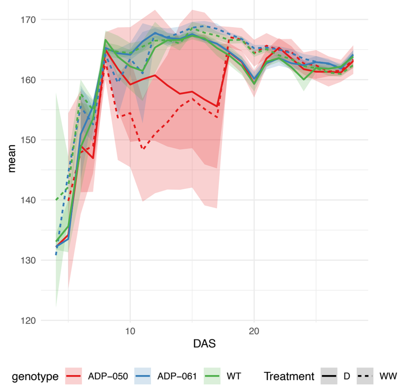 | | | 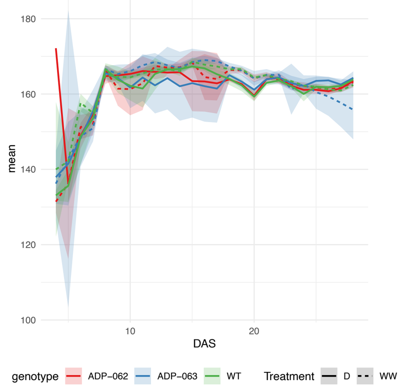 | | 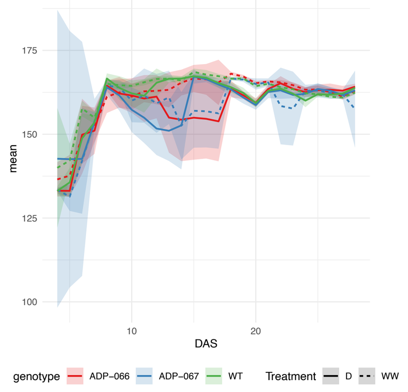 | | | | | 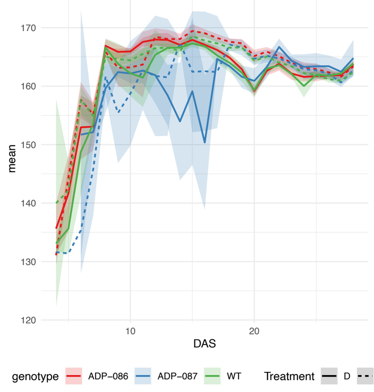 |
| top.intensity.fluo lab.b.mean | 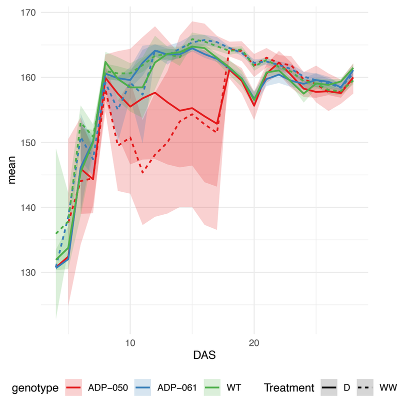 | | | | | 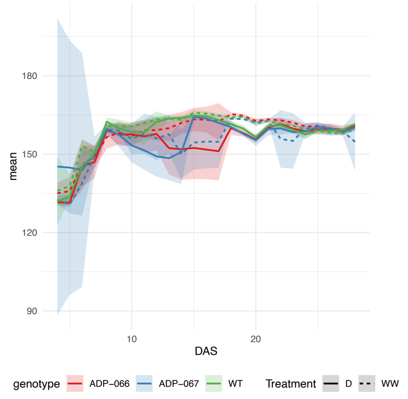 | | | | | 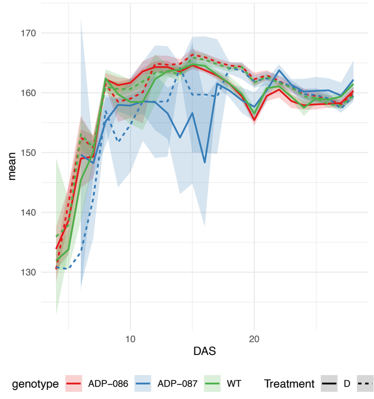 |
| top.intensity.fluo lab.l.mean | 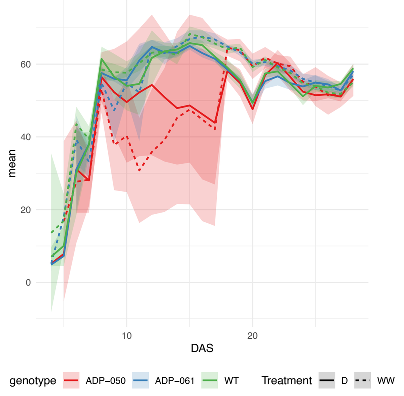 | | | | | 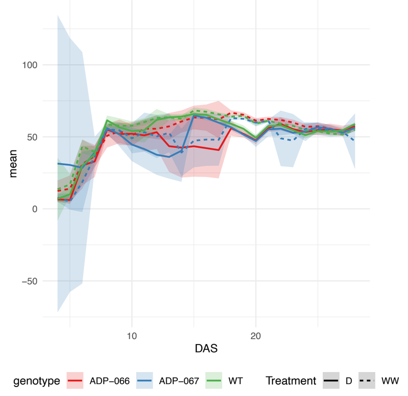 | | | | | 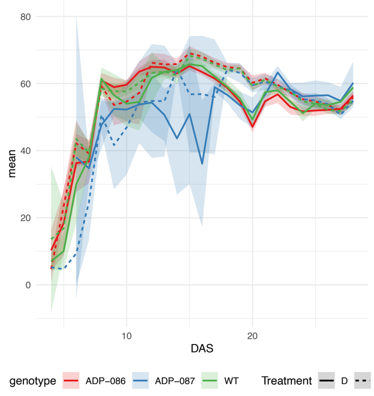 |
| top.intensity.nir mean | | 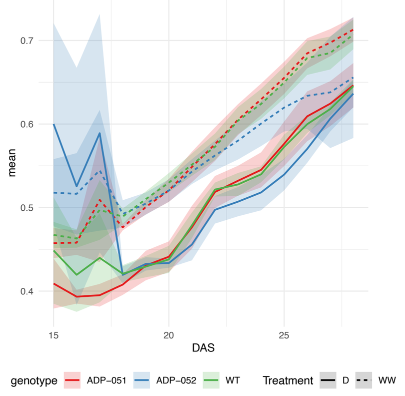 | | | | | | | | | |
| top.intensity.vis lab.b.mean | | | | | | | 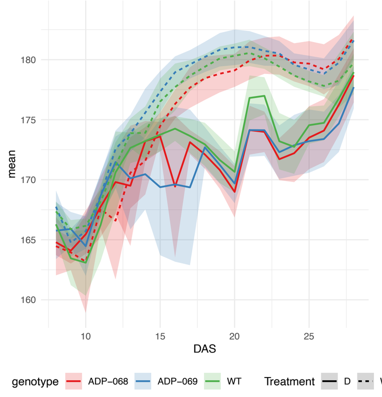 | 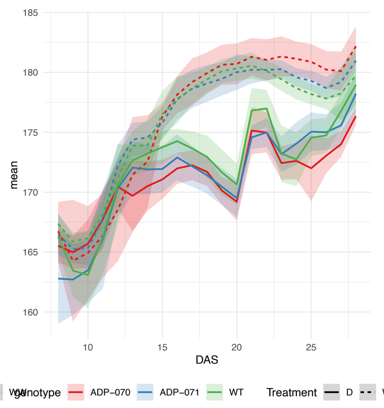 | 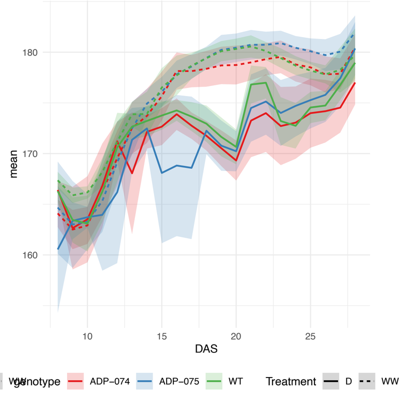 | | |
| top.intensity.vis rgb.blue.mean | | | 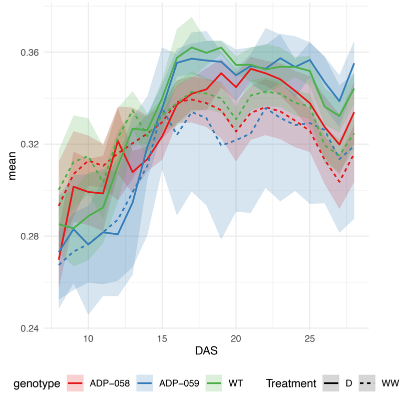 | | 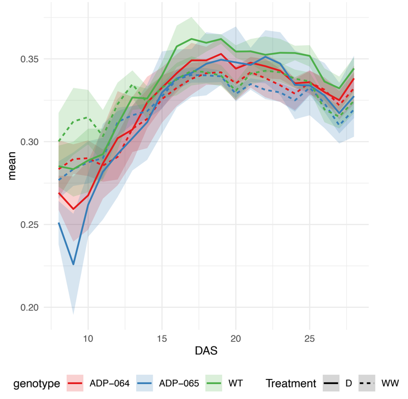 | | | | | 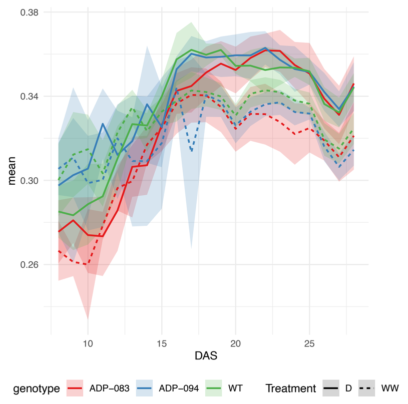 | |

| Genotype | Knockout.Line | Targeted.Gene | Primer.R | Primer.L |
|---|---|---|---|---|
| ADP−050 | SALK_023190C | AT3G51895 | AATTCCAAGAGAGGCTTCGAG | CTGGCTTTTGTGTCCTCTCTG |
| ADP−051 | GABI−KAT_571C05 | AT5G03260 | TAGGTACTCCAACGGTGTTTCTCT | GACCGTGGTGGAAATCGATG |
| ADP−052 | GABI−KAT_782F01 | AT5G03260 | TAGGTACTCCAACGGTGTTTCTCT | GTTTTAGTCAAAACCGACCGG |
| ADP−058 | GABI−KAT_760A01 | AT3G05630 | TCAGTATAGTGCAGAGGAAGAGCA | TCTTACACCTTGGAGCTTCAGT |
| ADP−059 | GABI−KAT_123E01 | AT3G05630 | TCCAAATGCAAACACTAAAAACTG | ACAGCAAGAAACGGCTTCAA |
| ADP−061 | SALK_060617C | AT3G51895 | AATGTTCCTATCCTGATGGGC | TGATGCTCCCATCTACTTTGC |
| ADP−062 | GABI−KAT_883C03 | AT5G54610 | TGTCCTGAAAGCATTAAAGATGTG | ATGTGAGATCCTCTGGCTCG |
| ADP−063 | GABI−KAT_537B07 | AT5G54610 | GTTTAAATGGTTGATCCGTGTTCT | TCAGGTCGCTTTTGGGTTTG |
| ADP−064 | SALK_118837C | AT3G06035 | TTCAGGATCTTTAGATCTGGTCTG | TCCGAGAATATCCTGCATTTG |
| ADP−065 | SALK_000844C | AT3G06035 | CTAGCTGACGAAATCGCAGAC | AGAGCCCAATTCGAGAGCTAC |
| ADP−066 | GABI−KAT_154B05 | AT3G45600 | ACAATTTAGGTCCTCCATGTGTTC | AGAAAATAAACAAAACAGAGCATTTCT |
| ADP−067 | GABI−KAT_026G04 | AT3G45600 | ACAATTTAGGTCCTCCATGTGTTC | GCAACAACCGGACTGCAAAT |
| ADP−068 | GABI−KAT_164E10 | AT3G16460 | TTTGACTTTTGATCGTTTAAGGCT | TGGTTGTGAAAGTCAACGCC |
| ADP−069 | GABI−KAT_146E08 | AT3G16460 | TTTGACTTTTGATCGTTTAAGGCT | GGTACCAAGGAAACCGGAGA |
| ADP−070 | GABI−KAT_306E08 | AT2G39350 | GAGATCCACTATACACGGTGTGG | ATCGGATTGCCAAAGGAAGC |
| ADP−071 | GABI−KAT_850E07 | AT2G39350 | CATGAAGATATAACGTTCCTGGAGA | CGTGTTCTCGGTTTGCTTGA |
| ADP−074 | GABI−KAT_161E04 | AT2G44450 | AATACTTGGTAGGTCGAAGGAGGT | AGCCTGGTTGATTCCTCCTT |
| ADP−075 | GABI−KAT_365C01 | AT2G44450 | TGAAAACCATGAGATTCTTCAAAA | GCAAATGTAGTGTTGGAGCCA |
| ADP−083 | GABI−KAT_920B04 | AT2G18450 | CGAGTTCCTAAAATGCAAAGTTTC | AGGTGATGATCCAGATGCTGT |
| ADP−086 | SALK_068648C | AT4G11070 | GGGGAAGCCTGTGTTAATCTC | GAAAGGTTCCAGGATCTCCAG |
| ADP−087 | SALK_105785C | AT4G11070 | GGGGAAGCCTGTGTTAATCTC | CAGGAAGATGTTGCCAAAGTG |
| ADP−094 | GABI−KAT_254B04 | AT2G18450 | CTTCTAAAATCCGCTTAAACATGC | TGGCCTATATTTACCCCAATGTG |
